# Emergence of the East-Central-South-African genotype of Chikungunya virus in Brazil and the city of Rio de Janeiro may have occurred years before surveillance detection

**DOI:** 10.1101/502443

**Authors:** Thiago Moreno L. Souza, Yasmine Rangel Vieira, Edson Delatorre, Giselle Barbosa-Lima, Raul Leal Faria Luiz, Alexandre Vizzoni, Komal Jain, Milene Mesquita, Nishit Bhuva, Jan F. Gogarten, James Ng, Riddhi Thakkar, Andrea Surrage Calheiros, Ana Paula Teixeira Monteiro, Patrícia T. Bozza, Fernando A. Bozza, Diogo A. Tschoeke, Luciana Leomil, Marcos Cesar Lima de Mendonça, Cintia Damasceno dos Santos Rodrigues, Maria C. Torres, Ana Maria Bispo de Filippis, Rita Maria Ribeiro Nogueira, Fabiano L. Thompson, Cristina Lemos, Betina Durovni, José Cerbino-Neto, Carlos M. Morel, W. Ian Lipkin, Nischay Mishra

**Affiliations:** Instituto Nacional de Infectologia (INI), Fundação Oswaldo Cruz (Fiocruz), Rio de Janeiro, RJ, Brazil; Center for Infection and Immunity, Mailman School of Public Health, Columbia University, New York, New York, USA; Laboratório de AIDS e imunologia Molecular, Instituto Oswaldo Cruz (IOC), Fiocruz, Rio de Janeiro, RJ, Brazil; Laboratório de Vírus Respiratório e do Sarampo, IOC, Fiocruz, Rio de Janeiro, RJ, Brazil; Laboratório de Imunofarmacologia, IOC, Fiocruz, Rio de Janeiro, RJ, Brazil; D’Or Institute for Research and Education (IDOR), Rio de Janeiro, RJ, Brazil; Instituto de Biologia, Universidade Federal do Rio de Janeiro (UFRJ), Rio de Janeiro, RJ, Brazil; Laboratório de Flavivírus, IOC, Fiocruz, Rio de Janeiro, RJ, Brazil; SAGE/COPPE, UFRJ, Rio de Janeiro, RJ, Brazil; Secretaria Municipal de Saúde, Rio de Janeiro, Brazil 34; National Institute for Science and Technology on Innovation on Diseases of Neglected Populations (INCT/IDPN), Center for Technological Development in Health (CDTS), Fiocruz, Rio de Janeiro, RJ, Brazil

**Keywords:** Chikungunya, arboviruses, surveillance, ECSA, Brazil, VirCapSeq-VERT

## Abstract

Brazil, which is hyperendemic for dengue virus (DENV), has had recent *Zika* (ZIKV) and (CHIKV) *Chikungunya* virus outbreaks. Since March 2016, CHIKV is the arbovirus infection most frequently diagnosed in Rio de Janeiro. In the analysis of 1835 syndromic patients, screened by real time RT-PCR, 56.4% of the cases were attributed to CHIKV, 29.6% to ZIKV, and 14.1% to DENV-4. Sequence analyses of CHIKV from sixteen samples revealed that the East-Central-South-African (ECSA) genotype of CHIKV has been circulating in Brazil since 2013 [95% bayesian credible interval (BCI): 03/2012-10/2013], almost a year before it was detected by arbovirus surveillance program. Brazilian cases are related to Central African Republic sequences from 1980’s. To the best of our knowledge, given the available sequence published here and elsewhere, the ECSA genotype was likely introduced to Rio de Janeiro early on 2014 (02/2014; BCI: 07/2013-08/2014) through a single event, after primary circulation in the Bahia state at the Northestern Brazil in the previous year. The observation that the ECSA genotype of CHIKV was circulating undetected underscores the need for improvements in molecular methods for viral surveillance.

## 1. Introduction

*Chikungunya virus* (CHIKV) is an alphavirus in the family *Togaviridae*, that frequently causes a febrile illness associated with arthralgia and skin rash, a classical triad of clinical manifestations classified as chikungunya fever^1^. Although severe prolonged and debilitating joint pain along with edema differentiate CHIKV infection from others caused by dengue (DENV) and Zika (ZIKV) for example – these arboviruses trigger similar symptoms, particularly during the early phase of infection^1^. Like ZIKV^2^, CHIKV infection has also been associated with the Guillain-Barre syndrome (GBS)^3^.

Chikungunya fever is a global public health problem with profound impact in tropical and subtropical regions of the world, wherein *Aedes (Stegomyia)* spp mosquitoes are especially prevalent and resources for mosquito abatement are limited^1^. There is no specific treatment or vaccine for CHIKV^4^; thus, vector control and avoidance are the main strategies currently available for disease control.

CHIKV has a positive-sense single-stranded RNA virus with a 11.8 kilobase genome that encodes two polyproteins, that are cleaved in four non-structural proteins (nsP1-ns4) and five structural proteins (C, E1, E2, E3 and 6K)^5^. Based on the genomic diversity of the CHIKV, or most often on the polymorphisms on the E1 region, different genotypes have been classified: East-Central-South-African (ECSA), West African and Asian. Adittionally, Indian Ocean lineage (IOL) appears to be emerging as an independent clade from the ECSA genotype^6^.

Since 2014, Asian and ECSA genotypes co-circulate in at North and Northeast regions of Brazil, respectively^7,8^, which raises the potential for co-infections and recombination. The ECSA genotype has been described in autochthonous cases in Rio de Janeiro^9–11^. Imported cases of Asian genotype have been described in Southeast Brazil^12^. The scale of the circulation of these different genotypes in Brazil is not known.

The Arbovirus Surveillance Program of the Municipal Health Department of Rio de Janeiro has recognized the city as historically hyperendemic for DENV, and since last years, both ZIKV and CHIKV were introduced. Here, after screening 1835 patients, we describe CHIKV ECSA genotype diversity and provide evidence of its introduction to Brazil in 2013 [95% Bayesian credible interval (BCI): 03/2012-10/2013], up to a year before surveillance detection. The mean time of ECSA introduction in Rio de Janeiro is on Febrary of 2014 (BCI: 07/2013-08/2014) in a single event, according to sequences available. Remarkably, the Brazilian cases are related to Central African sequences from 1980’s, highlighting that CHIKV ECSA circulation has been neglected for decades throughout the world.

## 2. Material and Methods

### 2.1. Study Population

Subjects were 1835 individuals with suspicious diagnosis of arbovirus infection, defined by fever (≥ 38 °C), exanthemata, headache, retro-orbital pain, photophobia, lumbar back pain, chills, weakness, malaise, nausea, vomiting or myalgia^1^, who presented within five days of illness onset to sentinel health care units or Quinta D’Or Hospital, a private and general hospital in the city of Rio de Janeiro, during the interval from March 2016 to June 2017. Samples were collected with informed consent in accordance with Institutional review board protocols approved by Fiocruz (# 57020616800005262 and 42999214110015248). Serum or plasma were tested for DENV, ZIKV and CHIKV; whereas urine was tested only for ZIKV.

### 2.2. RNA extraction

Viral RNA was extracted from serum or plasma and urine samples by QIAamp Viral RNA Mini Kit (Qiagen^®^, Dusseldorf, DE), eluted to a final volume of 60 µL and analyzed by performing real time RT-PCR assays. To evaluate cross-contamination, negative controls were handled at all stages. Procedures were conducted under biosafety level 2 or 3, according to international guidelines and Brazilian classification of pathogens^13,14^.

### 2.3. Real time RT-PCR

Amplification assays by real time RT-PCR were performed with QuantiTect/QuantiNova Probe RT-PCR Kit (Qiagen^®^) according to manufacturer’s conditions in 25 μL of reaction volume, including 5 μL RNA, 1μM each primers Forward/Reverse (F/R) and 0.2 μM probe shown in Table S1. Reverse transcription was carried out at 50 °C for 30 min, initial denaturation at 95 °C for 15 min, followed by 50 cycles of denaturation at 94 °C for 15s, and annealing at 55 °C for 35s. Samples with cycle threshold (ct) values lower than 40 and with sigmoid curves were considered positive.

Of note, the four serotypes of DENV, CHIKV and ZIKV were tested in the plasma or serum samples. The urine sample was tested for ZIKV RNA. For quality assurance purposes, different controls were included: mock-controls from extraction, non-template controls of RT-PCR reaction, positive controls for each virus and human 18S rRNA (ThermoFischer, Waltham, Masschusetts, USA) for each sample.

### 2.4. Sample election and Next Generation Sequencing (NGS)

Forty samples (serum or plasma), strongly positive for CHIKV RNA (as judged by ct values between 20 to 30) and negative for DENV and ZIKV, were further re-extracted and re-tested with CII-ArboViroPlex rRT-PCR assay^15^ to confirm molecular diagnosis. Subsequently, they were selected for Virome Capture Sequencing Platform for Vertebrate Viruses (VirCapSeq-VERT)^16^.

Total nucleic acid (TNA) was extracted from the ~200 μl volume of clinical sample using the easyMAG automated platform (Biomérieux^®^), following the manufacturer’s recommendations. Extracted TNA was eluted to a final volume of 50 µL in H^2^O. CII-ArboViroPlex rRT-PCR assay^15^ confirmed that samples are consistently positive for CHIKV and negative for ZIKV, all 4 serotypes of DENV and *West Nile* virus. Samples were additionally tested negative for *Mayaro* and *yellow fever* viruses, *Plasmodium spp*. and *Salmonella typhi* using in house multiplex assays. Of note, CII-ArboViroPlex rRT-PCR was more sensitive than the QuantiTect/QuantiNova Probe RT-PCR Kit in detecting CHIKV (Figure S1).

Fourteen samples, with the lowest ct values, distributed temporally across the sampling period were enriched using the VirCapSeq-VERT protocol^16^ (Table S2). Sequencing was performed on the Illumina MiSeq platform (Illumina, San Diego, CA, USA) with Reagent kit v3 resulting in 30,393,722 paired end (300 bp) reads. Two additional 2015 samples were submitted for unbiased sequencing following Ribo-zero treatment to deplete ribosomal-RNA sequences, as previously described^17^. More specific details on sequencing protocols and analysis are described under the Supplemental Methods. Genome recovery was greater than 99%, except for one sample with 95% genome recovery (Table S2).

### 2.5. Phylogenetic analysis

To investigate the origin of CHIKV in Rio de Janeiro, the newly generated genomes obtained via VirCapSeq and unbiased sequencing were included in an analysis with all CHIKV genomes deposited in GenBank before October 2017 with more than 10,000 nt^18^. This resulted in a final dataset of 555 genomic sequences representing all three viral genotypes and the IOL clade (Table S3). Non-aligned terminal sequences were trimmed before analyses. After sequence alignment using MAFFT^19^, viral phylogenies were reconstructed by maximum likelihood (ML) analysis implemented in RaxML^20^. The pattern of spatiotemporal viral diffusion and the ancestral complete coding sequences (CDS) at key internal nodes of the ECSA genotype phylogeny were reconstructed only for the ECSA genotype (excluding IOL strains, Table S3) by Bayesian inference with Markov chain Monte Carlo (MCMC) sampling as implemented in BEAST v1.8 package^21^ using the GTR+Γ^4^ nucleotide substitution model selected by jModelTest v1.6^22^. We removed from the ECSA genotype dataset sequences wherein more than 20% of bases were indeterminate, those that lacked geographical and temporal information or had been passaged multiple times in culture. Before the phylogeographic analysis, the temporal signal of the sequence dataset was tested with Tempest^23^. Comparisons between multiple combinations of non-parametric demographic models (skyline^24^ and skygrid^25^), molecular clock models (strict and relaxed uncorrelated molecular clock [UCLN] models^26^), and reversible (symmetric) and nonreversible (asymmetric) discrete phylogeographic models^27^ were performed using the log marginal likelihood estimation (MLE) based on path sampling (PS) and stepping-stone sampling (SS) methods^28^. Analyses were run for 10^8^ generations and convergence (effective sample size > 200) was inspected using TRACER v1.6 (http://tree.bio.ed.ac.uk) after discarding 10 % burn-in. The maximum clade credibility tree was summarized using TreeAnnotator v.1.8^21^ and visualized with FigTree v.1.4.2 (http://tree.bio.ed.ac.uk). Ancestral complete coding sequences at key internal nodes of the ECSA phylogeny were reconstructed in BEAST v1.8 package and synonymous and nonsynonymous substitutions were annotated using the Geneious v9 program.

## 3. Results

### 3.1. Since March 2016, CHIKV is the most common arbovirus in Rio de Janeiro, Brazil

A total of 1835 patients in Rio de Janeiro with arbovirus-like illness were tested by real time RT-PCR for the presence of CHIKV, DENV (all four serotypes) and ZIKV RNA from March 2016 to June 2017 (Figure 1). Approximately 70 % of these patients presented during the summer months (end of December to end of March) (Figure 1) coincident with higher mosquito population levels. Socio-demographical data was available for 99.5 % of these patients (Table S4), revealing that most of them were young adult females. The frequency of arboviruses detection was higher in 2016 than in 2017 (Figure 1 and Table S4). In 2016, CHIKV, DENV-4 and ZIKV RNA was found in 72, 12 and 16% of the patients, respectively. During the studied period of 2017, CHIKV, DENV-4 and ZIKV RNA was found in 37, 17 and 47% of those with positive results, respectively.

**Figure 1.**
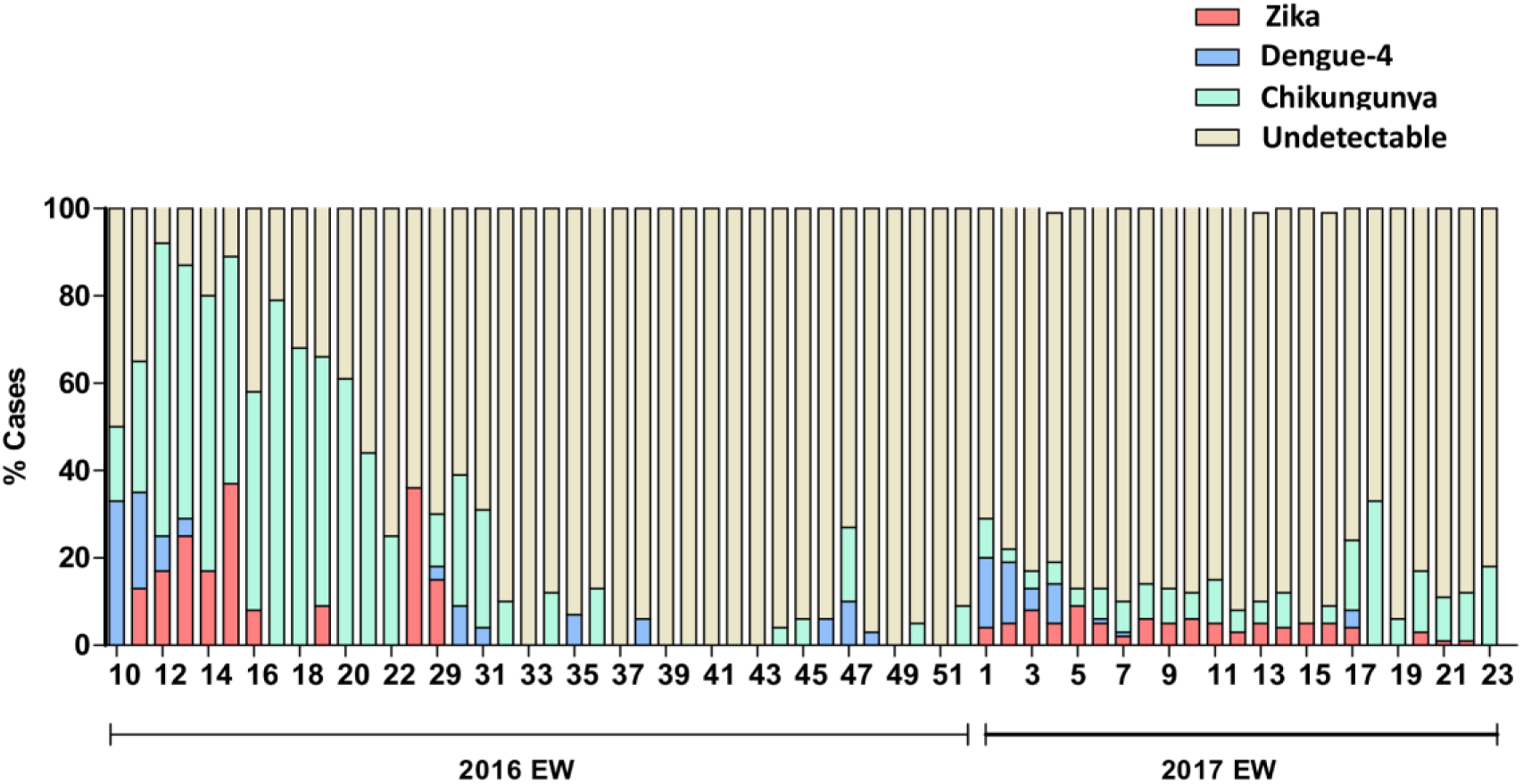
Molecular positivity for arboviruses in the city of Rio de Janeiro, Brazil, from March 2016 to June 2017. Percentage of positive cases (y-axis) for CHIKV, ZIKV and DENV-4, and the negative ones are indicated by each epidemiological week (x-axis).

The cases of CHIKV, DENV-4, and ZIKV also overlapped geographically within the city (Table S4, Figure S2 and S3), highlighting the wide spread circulation of these arboviruses. Altogether the majority of the confirmed cases throughout the studied period was related to CHIKV infection (Figure 1 and Table S4).

### 3.2. Phylogenetic analysis of CHIKV

The ML phylogenetic analysis indicated that all Brazilian sequences from Rio de Janeiro belonged to the ECSA genotype with limited genomic diversity among strains, forming a statistically supported cluster (CHIKV-RJ, bootstrap = 79%) within a monophyletic clade (bootstrap = 100%) comprising all other Brazilian ECSA sequences (Figure 2). The temporal analysis of the ECSA sequences showed a strong correlation (*R*^2^ = 0.97) between genetic divergence and sampling time (Figure 3), supporting the use of temporal calibration directly from sequences.

**Figure 2.**
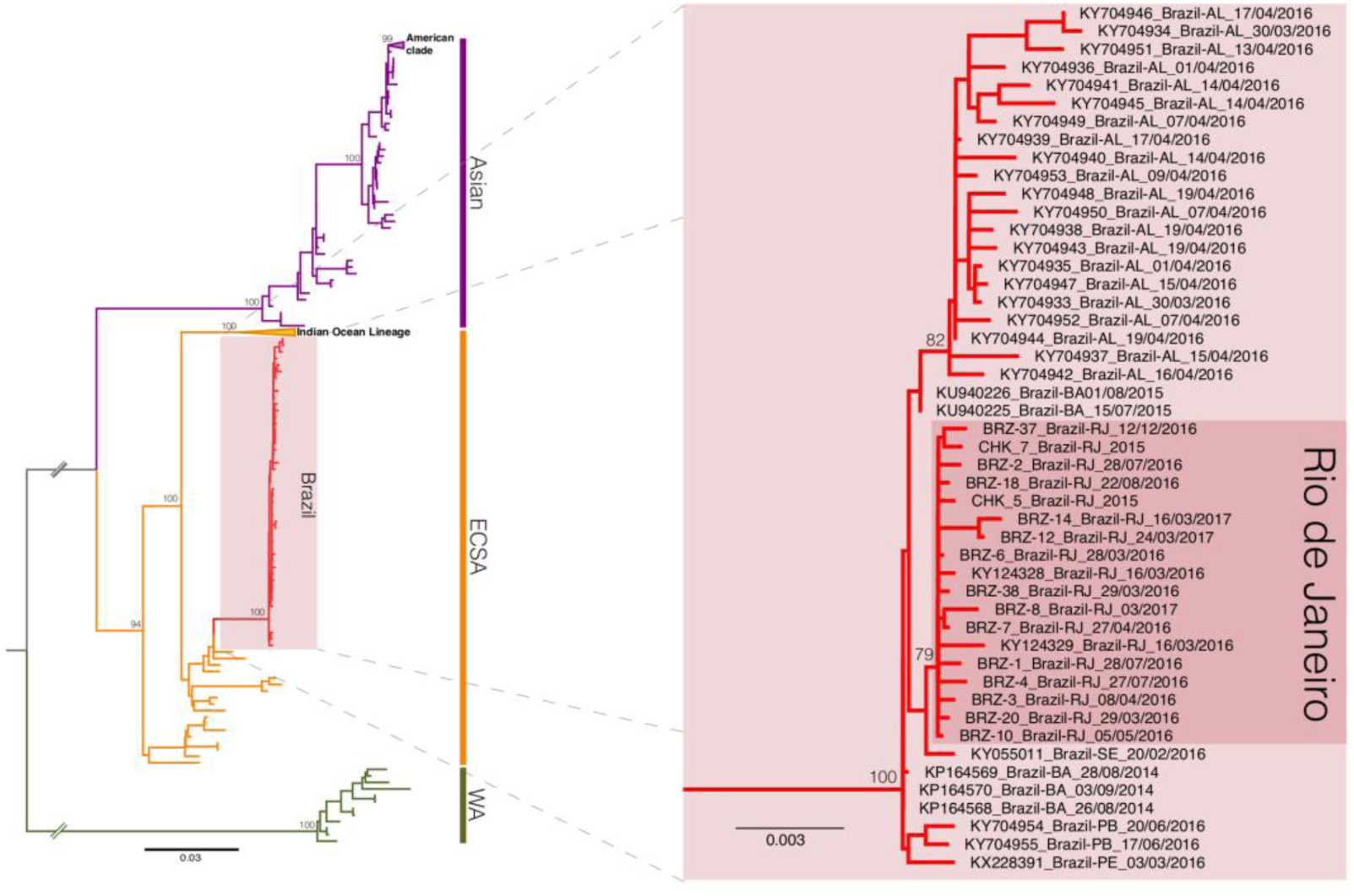
Maximum likelihood phylogeny of the CHIKV full-length genome dataset. The bootstrap values are indicated for each genotype-specific clade (vertical bars, ECSA: East-Central-South African, WA: West African) and important intra-genotype lineages (Asian: American and ECSA: Indian Ocean Lineage and Brazil). The inset offer a close view of the ECSA genotype clade showing the Brazilian cluster (pink box) and the inner Rio de Janeiro cluster (red box). The branch lengths are drawn to scale with bar at the bottom indicating nucleotide substitutions per site.

**Figure 3.**
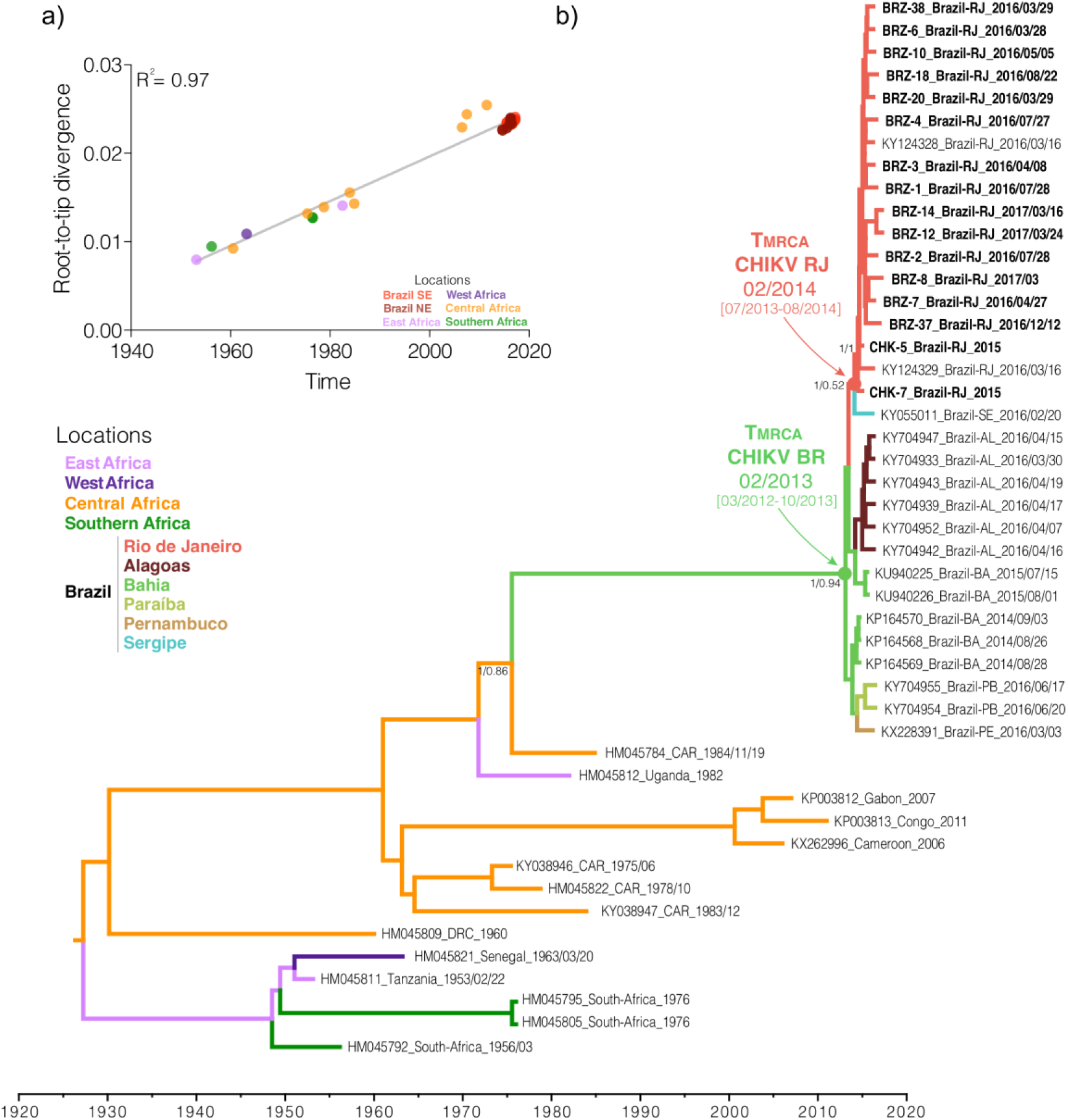
Phylogeography of the CHIKV ECSA genotype. a) Temporal signal analysis correlating the sampling date of each sequence and its genetic distance from the root of a maximum likelihood phylogeny (R^2^ = 0.97). b) Time-scaled Bayesian phylogeographic MCC tree of the CHIKV ECSA genotype full-length genomes. The colors of branches represent the most probable location of their descendent nodes as indicated at the legend (bottom right). Branch support are indicated only at key nodes (posterior/posterior state probability). The nodes representing the ECSA introduction in Brazil (red dot) and Rio de Janeiro (green dot) are indicated. All horizontal branch lengths are drawn to a scale of years. Tips names were coded as accession number_country_date. (AL: Alagoas state, BA: Bahia state, PA: Paraiba state, PE: Pernambuco state, RJ: Rio de Janeiro state, SE: Sergipe state; CAR: Central African Republic; DRC: Democratic Republic of Congo; USA: United States of America).

The evolutionary rate of the ECSA genotype estimated in this study was 2.85 × 10^-4^ substitutions/site/year (BCI: 2.50 – 3.23 × 10^-4^ substitutions/site/year), and was consistent among the different molecular clock and coalescent models evaluated (Figure S4 and Table S5). The ECSA genotype evolutionary rate found in this study was similar to previous estimates for the ECSA and other CHIKV genotypes^8,29^. The time-scaled phylogenetic tree estimated the introduction of the ECSA genotype in the State of Bahia, Northeastern Brazil, to the beginning of 2013 [(BCI: March 2012 - October 2013] probably from a Central African country (posterior state probability, *PSP* = 0.86) with spread thereafter to other Brazilian states in the Northeastern (Alagoas, Paraíba and Pernambuco, *PSPs* ≥ 0.54) and Southeastern (Rio de Janeiro, *PSPs* = 0.52) regions. The introduction of the ECSA genotype to Rio de Janeiro was dated to early 2014 (mean time Febrary 2014; BCI: July 2013 – August 2014). From Rio de Janeiro, this lineage returned to Northeast region, spreading to Sergipe. However, the low *PSP* sustaining this viral flux does not allow us to exclude the possibility that this node was in Bahia (*PSP* = 0.35). In this alternative scenario, the introduction of the ECSA genotype in Rio de Janeiro might have occurred in middle 2014 (BCI: February 2014 – October 2014).

No substitution in the envelope proteins E1 (A226V, K211E) and E2 (L210Q, V264A) associated with enhanced CHIKV fitness in *Ae. aegypti* and *Ae. albopictus*^30^ were detected in the sequences from Rio de Janeiro. Ancestral genomic sequences reconstruction showed that 16 amino acid substitutions were fixed between the divergence of the CHIKV-BR clade and its MRCA in Central Africa (Table S6). These substitutions displayed a balanced distribution between nonstructural (*n* = 7) and structural (*n* = 9) proteins. The ancestral inference also revealed that four amino acid substitutions accumulated in the ECSA lineage introduced in Rio de Janeiro. The change P352A in nsP2 seems to have emerged in Bahia state, and spread to Alagoas state. According to our phylogeographic reconstruction, the remaining three amino acid mutations arose after the introduction of the ECSA genotype in Rio de Janeiro. Two of these changes, one in E1 (K211T) and one in nsP4 (A481D), spread to Sergipe. The I111V substitution in nsP4 was found only in isolates from Rio de Janeiro.

## Discussion

Recent studies suggest that ZIKV circulated in the Americas for several months prior to detection^31,32^. Our findings demonstrate an similar scenario for CHIKV, wherein the virus may have circulated for up to one year before its detection. During the period in which surveillance was increased in Rio de Janeiro due to ZIKV concerns, CHIKV has become the most prevalent arbovirus in the city. Surveillance data reveal majority of screened individuals had undetectable viral loads for DENV, ZIKV or CHIKV. Most likely, these results are due to a broad/inclusive case definition (fever or exanthema plus another symptom) used by the surveillance system, which could be caused by various infectious and non-infectious conditions. Nevertheless, unbiased molecular diagnostic tools could be worthwhile to detect if other pathogens could be also circulating in the city. These observations support the use of multiplexed and unbiased diagnostic assays in public health surveillance, in order to extend the diagnostic coverage.

Based on the notification of the Brazilian surveillance systems, CHIKV was introduced in Brazil during 2014^7,8^, the Asian genotype was confirmed in the North Region of Brazil (Oiapoque, Amapá state) and the ECSA genotype was first identified in the Northeastern region of Brazil (Feira de Santana, Bahia state). The ECSA genotype was subsequently detected in other states, particularly Sergipe^33^ and Rio de Janeiro^11^, on the Northeastern and Southeastern regions, respectively. Remarkably, the first documented cases appeared in Rio de Janeiro in late 2015^34^. The ECSA genotype now appears to be well established in the city of Rio de Janeiro, with a clear genetic signature (I111V in nsP4). Confidence intervals for the timing of the arrival of CHIKV in Brazil and Rio de Janeiro range from 2012 to 2014, the entire time interval suggests the virus may have been present for some time before surveillance detection^7,8,12^.

Our results through phylogenomic analyses suggest that CHIKV ECSA genotype was likely introduced in a single event into Brazil in 2013, up to one year before previous estimates^35^. Previous finding correlate Brazilian ECSA genotypes with an Angola strain from 1962^35^. We removed the sequence from Angola-1962 from our dataset due to multiple passages in cell culture^36^, which may have introduced cell-derived genetic drifts in the virus genome. Our phylogeographic reconstruction suggests that the Central African region is the probable source of the ECSA lineage that spread to Brazil. However, sufficient data on ECSA genomes from Central Africa is not available for a definitive conclusion. Thus, stronger sampling of CHIKV strains at that region could increase the molecular epidemiology understanding of the overlooked CHIKV ECSA genotype circulation from the 1980’s to contemporary, helping to point out more precisely the country of origin of the Brazilian ECSA outbreak.

We have analyzed sequences from seven Brazilian states (Alagoas, Bahia, Sergipe, Paraíba, Pernambuco and Rio de Janeiro) from two regions, spanning a four-year time interval (2014–2017). Our results corroborate that the introduction of the ECSA genotype in Brazil most probably occurred in Bahia^35^; however, our analysis, differs in the mean time of the introduction of the ECSA genotype up to one year before previous estimates^35^. This discrepancy in the estimates of the date of the most recent common ancestor of the Brazilian ECSA lineages may reflect access to higher number of sequences used in the present study that enhance the accuracy of modeling. From Bahia state, the ECSA genotype spread to other Northeastern states (Alagoas and Paraiba) and to Rio de Janeiro (Southeast region). Our temporal reconstruction indicates that the introduction of the ECSA genotype in Rio de Janeiro most probably occurred in early 2014 and that from Rio de Janeiro, this lineage returned to Northeastern region, spreading to Sergipe.

In addition to the E1 K211T substitution previously described as a genetic signature of the CHIKV isolates from Rio de Janeiro^11^, we found three additional amino acid mutations in the nonstructural proteins nsP2 (P352A) and nsP4 (I111V and A481D). The I111V substitution nsP4 was exclusively found in CHIKV ECSA strains from Rio de Janeiro. The effects of these substitutions remains to be elucidated; however, it is noteworthy that the E1 K211T substitution is positioned at the same site of the substitution K211E, previously associated with enhanced fitness in *Aedes aegypti*, when in an E1-226A background.

Our work highlights that CHIKV became the most prevalent arbovirus in the city of Rio de Janeiro in March 2016. Sequenced CHIKV samples revealed the presence of the ECSA genotype, which is likely circulating Brazil for up to one year before detection by surveillance systems. Of note, both un- and biased sequencing technologies were used in this study. VirCapSeq^16^ enhanced our sequencing capacity over 600-times in comparison to unbiased high throughput sequencing, with respect to the average depth per bp.

Altogether, we showed here a consistant CHIKV activity in Rio de Janeiro since 2016, a probable introduction of the ECSA genotype in Brazil and Rio de Janeiro up to a year earlier than previously thought. Periods of cryptic transmission, desmonstrated here for CHIKV, reinforce the importance of the continuous surveillance activity along with genomic data to provide timely information to orientate effective public health responses.

## AUTHOR CONTRIBUTIONS

Laboratorial-based Surveillance – TMLS, YRV, GB-L, RLFL, AV, JN, RT, AMBF, RMRN

Patients Enrolment – PTB, FAB, ASC, APTM

Clinical Surveillance – CL, BD, JC-N

Sequencing - NB, JFG, DAT, LL, MCLM, CDSR, MCT, FLT, MM

Bioinformatics – KJ, ED

Study coordination – TMLS, CMM, WIL, NM

Manuscript preparation and revision – TMLS, YRV, ED, WIL, NM

All authors revised and approved the manuscript.

TMLS, YRV and ED – contributed equally as first author

WIL and NM – contributed equally as last author

## Financial Support

This study was financed in part by the Coordenação de Aperfeiçoamento de Pessoal de Nível Superior - Brasil (CAPES) - Finance Code 001. YRV and ED are funded by fellowships from the Coordenação de Aperfeiçoamento de Pessoal de Nível Superior - Brasil (CAPES). Funding was also provided by National Council for Scientific and Technological Development (CNPq), Ministry of Science, Technology, Information and Communications (no. 465313/2014-0); Ministry of Education/CAPES (no. 465313/2014-0); Research Foundation of the State of Rio de Janeiro/FAPERJ (no. 465313/2014-0) and Oswaldo Cruz Foundation/FIOCRUZ. The study was also supported by the National Institutes of Health (U19 AI109761) to NM, WIL, KJ, RT, JN, NB at the Center for Infection and Immunity at Columbia University, New York; and JFG is a fellow working at CII who was supported by the DAAD with funds from the German Federal Ministry of Education and Research (BMBF) and the People Programme (Marie Curie Actions) of the European Union’s Seventh Framework Programme (FP7/2007-2013) under REA grant agreement n° 605728 (P.R.I.M.E. – Postdoctoral Researchers International Mobility Experience). Funders had no role in the experiment design or interpretation.

## Acknowledgements

We appreciate the contribution from Rio de Janeiro’s city surveillance SMS-RJ.

## Conflict of Interest

None declared.

