## Supplemental Material for "Emergence of the East-Central-South-African genotype of Chikungunya virus in Brazil and the city of Rio de Janeiro may have occurred years before surveillance detection"

**Affiliations:**

*Supplemental Methodology*

**Downstream processing of the Full genome sequencing using VirCapSeq-VERT platform**

Sequencing on the Illumina MiSeq platform (Illumina, San Diego, CA, USA) resulted in 30,393,722 (300 bp) paired end reads. The demultiplexed FastQ files were adapter trimmed using cutadapt program (v 1.8.3). Adaptor trimming was followed by generation of quality reports using FastQC software (v 0.11.5) which were used to determine filtering criteria based on average quality scores of the reads, presence of indeterminate nucleotides and homopolymeric reads. The reads were quality filtered and end-trimmed with PRINSEQ software (v 0.20.3). The filtered fastq files were mapped to the complete genome sequence of Chikungunya virus using Bowtie 2 mapper (v 2.2.9). The consensus genome sequences from mapping assemblies were obtained using SAM Tools (v 1.3.1) and Bcftools (v 1.3.1). The genome variation was analyzed using in-house perl scripts to count the frequency of nucleotides at each position with reference to the Chikungunya genome.

**Read processing**

The demultiplexed FastQ files were adapter trimmed using cutadapt program (v 1.8.3)[5]. Adaptor trimming was followed by generation of quality reports using FastQC software (v 0.11.5) which were used to determine filtering criteria based on average quality scores of the reads, presence of indeterminate nucleotides and homopolymeric reads[6]. Reads were quality filtered and end-trimmed with PRINSEQ software (v 0.20.3)[7]. The filtered fastq files were mapped to the complete genome sequence of CHIKV using Bowtie2 mapper (v 2.2.9)[8]. The consensus genome sequences from mapping assemblies were obtained using SAM Tools (v 1.3.1) and Bcftools (v 1.3.1)[9]. Genome variation was analyzed using in-house perl scripts to count the frequency of nucleotides at each position with reference to the Chikungunya genome.

**Unbiased-NGS**

Additionally, two further CHIKV positive samples from 2015 were sequenced by unbiased shotgun sequencing. In brief, 0.3 mL plasma aliquot was extracted using QIAamp Viral RNA Mini Kit (Qiagen®) with RNase-free DNase (Qiagen®) treatment, omitting carrier RNA. Double-stranded cDNA libraries were constructed using a TruSeq Stranded Total RNA LT kit (Illumina®) with Ribo-zero treatment. The library size distribution was assessed using a 2100 Bioanalyzer (Agilent®) with a High Sensitivity DNA kit (Agilent®), and the quantification was performed using a 7500 Real-time PCR System (Applied Biosystems®) with a KAPA Library Quantification Kit (Kapa Biosystems). Paired-end sequencing (2 x 300 bp) was performed with a MiSeq Reagent kit v3 (Illumina®). The sequences obtained were preprocessed using the PRINSEQ software to remove reads smaller than 50 bp and sequences with scores of lower quality than a Phred quality score of 20. Paired-End reAd merger (PEAR) software was used to merge and extend the paired-end Illumina reads using the default parameter. The extended reads were analyzed against the Human Genome Database using the DeconSeq program, with an identity and coverage cutoff of 70%, to remove human RNA sequences. Non-human reads were analyzed against all GenBank viral genomes (65,052 sequences) using the BLAST software with a 1e-5 e-value cutoff. The sequences rendering a genome were assembled with SPAdes 3.7.1 software[10] followed by a second assembly with the CAP3 program using default parameters[11].

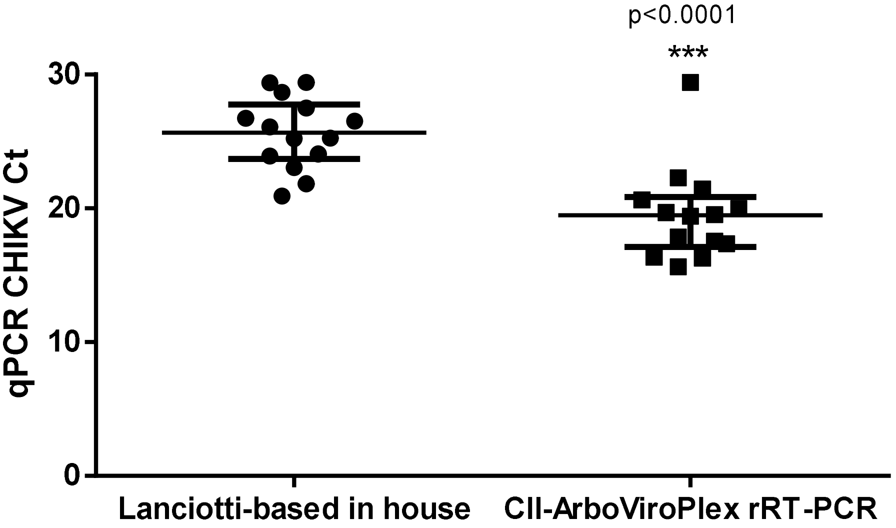

**Figure S1. Comparison of diagnostic sensitivities among different acid nucleic extraction and RT-PCR methodologies.** CHIKV cycle threshold (Ct) values obtained for 14 RNA samples collected between Feb 2016 - Feb 2017 processed by column-based extraction methods and qPCR *in house*[1–3] ranged from 20.92 to 29.42. Ct values obtained for the same samples after extraction methods based on magnetic beads and the new 5-ArboViroPlex qPCR[4] ranged from 16.3-29.41. Statistical analysis to compare sensitivities of CHIKV molecular diagnostics was performed in accordance with paired t test (95% confidence intervals). A p-value < 0.05 was considered statistically significant. CHIKV viral load values were significantly higher by using an automated purification method and the 5-ArboViroPlex qPCR (p < 0.0001).

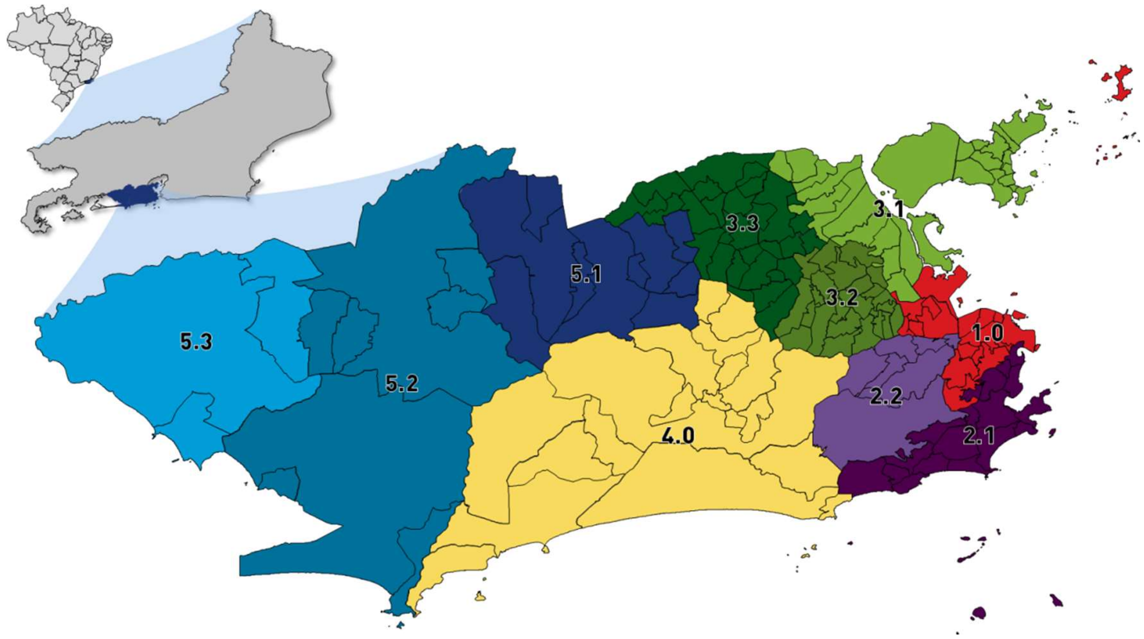

**Figure S2.** The City of Rio de Janeiro, Brazil and its planning areas (AP), distributed according to municipal administration.

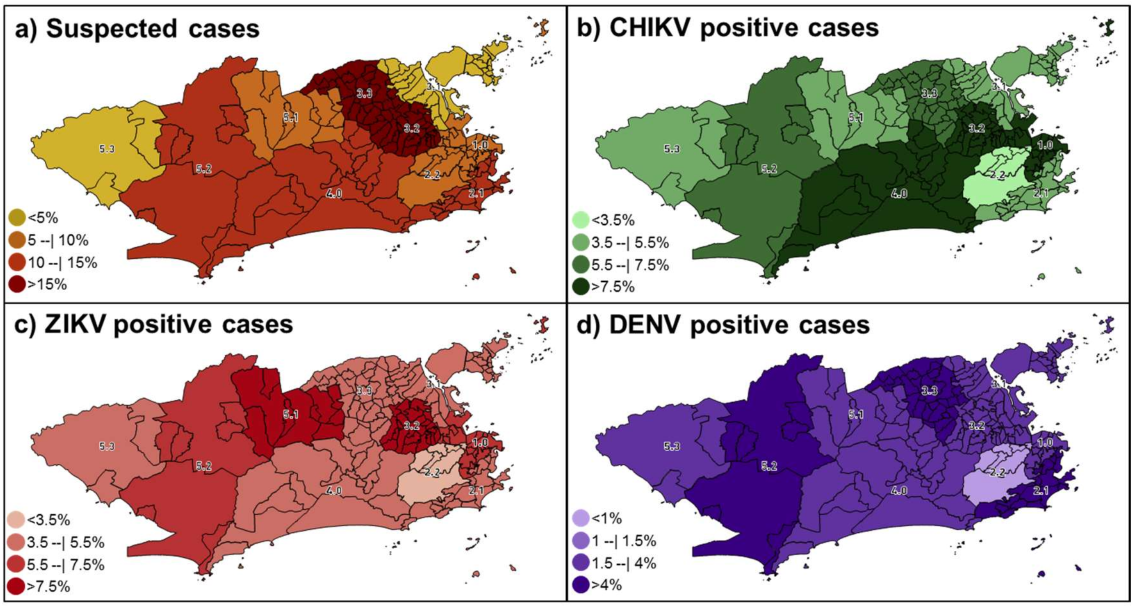

**Figure S3. Geographic distribution of suspected and positive cases for arboviruses in the city of Rio de Janeiro, Brazil, from March 2016 to June 2017.** Percentage of suspected (a) and positive cases for CHIKV (b), ZIKV (c) and DENV-4 (d) by PA. For guidance, the site where the patient was medically assisted was taken as reference. These percentages are not absolute indications that a certain PA is more affected by one virus than another. Because individuals may live in PA, work and seek for medical assistance in other regions. These data robustly show that cases of either DENV, ZIKV and CHIKV are widespread throughout the city.

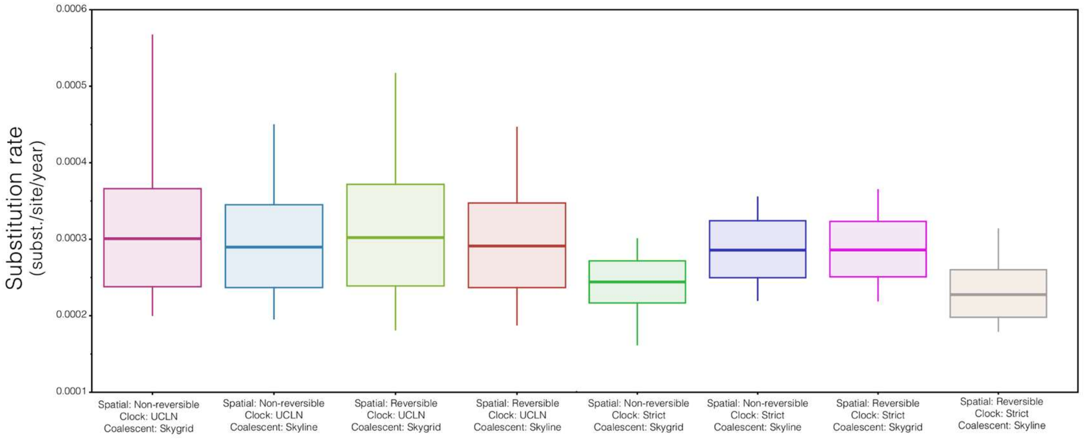

**Figure S4. BEAST estimates of the CHIKV ECSA substitution rates employing different model combinations.** The box plots represent the median substitution rates (substitutions/site/year) and the 95% Bayesian credible intervals of the posterior distributions estimated under each model combination as informed in the x-axis.

***Table S1.*** *Description of published primer/probe sets used in qPCR.*

| Virus Target | Reference | Primer/Probe | Genome Position | Sequence (5´- 3´) |
|---|---|---|---|---|
| ZIKV | Lanciotti *et al*, 2008[1] | Primer F | 1086-1102 | CCG CTG CCC AAC ACA AG |
| | | Primer R | 1162-1139 | CCA CTA ACG TTC TTT TGC AGA CAT |
| | | Probe | 1107-1137 | FAM/ AGC CTA CCT TGA CAA GCA GTC AGA CAC TCA A /BHQ1 |
| CHIKV | Lanciotti *et al*, 2007[2] | Primer F | 874-894 | AAA GGG CAA ACT CAG CTT CAC |
| | | Primer R | 961-942 | GCC TGG GCT CAT CGT TAT TC |
| | | Probe | 899-923 | FAM/ CGC TGT GAT ACA GTG GTT TCG TGT G/BHQ1 |
| DENV1 | Santiago *et al*, 2013[3] | Primer F | 8936-8955 | CAA AAG GAA GTC GYG CAA TA |
| | | Primer R | 9023-9047 | CTG AGT GAA TTC TCT CTG CTR AAC |
| | | Probe | 8961-8979 | FAM/CAT GTG GYT GGG AGC RCG C/BHQ1 |
| DENV2 | Santiago *et al*, 2013[3] | Primer F | 1426-1447 | CAG GCT ATG GCA CYG TCA CGA T |
| | | Primer R | 1482-1504 | CCA TYT GCA GCA RCA CCA TCT C |
| | | Probe | 1454-1480 | VIC/CTC YCC RAG AAC GGG CCT CGA CTT CAA/BHQ1 |
| DENV3 | Santiago *et al*, 2013[3] | Primer F | 701-720 | GGA CTR GAC ACA CGC ACC CA |
| | | Primer R | 749-775 | CAT GTC TCT ACC TTC TCG ACT TGY CT |
| | | Probe | 722-747 | FAM/ACC TGG ATG TCG GCT GAA GGA GCY TG/BHQ1 |
| DENV4 | Santiago *et al*, 2013[3] | Primer F | 884-904 | TTG TCC TAA TGA TGC TRG TCG |
| | | Primer R | 953-973 | TCC ACC YGA GAC TCC TTC CA |
| | | Probe | 939-965 | FAM/TYC CTA CYC CTA CGC ATC GCA TTC CG/BHQ1 |

**Bibliographical references are indicated at the end of this file*

**Table S2 - Quality control of genome assembly**

| Sample Name | GenBank ID | Collection date | Collection site* | Lanciotti-based in house Cts | CII Arboviro-Plex Cts | Sequence technology | Number rawfastq reads | Number input reads | Number mapped reads | Mapped reads (%) | Genome length | CHIKV used for mapping reference | Refseq Length | Number Mapped Pos Ref (Length Genome Recovered) | Genome Recovered (%) | Avg. depth/bp | Avg. unmapped length | Longest Unmapped Start | Longest Unmapped End |
|---|---|---|---|---|---|---|---|---|---|---|---|---|---|---|---|---|---|---|---|
| BRZ-10_S8_L001_R1_001 | MG649972 | May 04, 2016 | PA 1.0 | 26.08 | 16.35 | ViralCapSeq-VERT | 2,71E+06 | 2,65E+06 | 7,86E+05 | 29.71 | 1,17E+04 | NC_004162.2 | 1,18E+04 | 1,18E+04 | 99.72 | 9299 | 33 | 1,18E+04 | 1,18E+04 |
| BRZ-12_S9_L001_R1_001 | MG649982 | Mar 19, 2017 | PA 5.2 | 26.50 | 19.54 | ViralCapSeq-VERT | 2,27E+06 | 2,19E+06 | 1,01E+06 | 46.26 | 1,28E+04 | NC_004162.2 | 1,18E+04 | 1,18E+04 | 100 | 12082 | NA | NA | NA |
| BRZ-14_S10_L001_R1_001 | MG649978 | Mar 14, 2017 | PA 5.2 | 26.74 | 19.68 | ViralCapSeq-VERT | 2,51E+06 | 2,41E+06 | 2,54E+05 | 10.53 | 1,17E+04 | NC_004162.2 | 1,18E+04 | 1,18E+04 | 99.81 | 3104 | 22 | 1,18E+04 | 1,18E+04 |
| BRZ-18_S11_L001_R1_001 | MG649971 | Aug 21, 2016 | PA 3.3 | 27.48 | 21.44 | ViralCapSeq-VERT | 1,39E+06 | 1,31E+06 | 1,69E+04 | 1.28 | 1,17E+04 | NC_004162.2 | 1,18E+04 | 1,18E+04 | 99.66 | 202 | 40 | 1,18E+04 | 1,18E+04 |
| BRZ-1_S1_L001_R1_001 | MG649977 | July 27, 2016 | PA 5.2 | 20.92 | 17.55 | ViralCapSeq-VERT | 2,27E+06 | 2,20E+06 | 2,54E+05 | 11.51 | 1,17E+04 | NC_004162.2 | 1,18E+04 | 1,18E+04 | 99.79 | 3007 | 24 | 1,18E+04 | 1,18E+04 |
| BRZ-20_S12_L001_R1_001 | MG649976 | Mar 28, 2016 | PA 1.0 | 28.66 | 19.98 | ViralCapSeq-VERT | 2,03E+06 | 1,96E+06 | 7,80E+04 | 3.99 | 1,17E+04 | NC_004162.2 | 1,18E+04 | 1,18E+04 | 99.75 | 906 | 29 | 1,18E+04 | 1,18E+04 |
| BRZ-2_S2_L001_R1_001 | MG649981 | Jul 27, 2016 | PA 3.2 | 21.84 | 17.85 | ViralCapSeq-VERT | 1,50E+06 | 1,44E+06 | 8,19E+04 | 5.69 | 1,17E+04 | NC_004162.2 | 1,18E+04 | 1,18E+04 | 99.96 | 982 | 4 | 1,18E+04 | 1,18E+04 |
| BRZ-37_S13_L001_R1_001 | MG649973 | Dec 12, 2016 | PA 4.0 | 29.40 | 22.30 | ViralCapSeq-VERT | 1,88E+06 | 1,82E+06 | 1,21E+05 | 6.63 | 1,17E+04 | NC_004162.2 | 1,18E+04 | 1,18E+04 | 99.72 | 1363 | 32 | 1,18E+04 | 1,18E+04 |
| BRZ-38_S14_L001_R1_001 | MG649970 | Mar 29, 2016 | PA 1.0 | 29.42 | 20.63 | ViralCapSeq-VERT | 1,21E+06 | 1,17E+06 | 4,01E+04 | 3.44 | 1,17E+04 | NC_004162.2 | 1,18E+04 | 1,18E+04 | 99.56 | 477 | 51 | 1,18E+04 | 1,18E+04 |
| BRZ-3_S3_L001_R1_001 | MG649974 | Apr 07, 2016 | PA 1.0 | 23.04 | 15.63 | ViralCapSeq-VERT | 3,66E+06 | 3,59E+06 | 4,00E+05 | 11.13 | 1,17E+04 | NC_004162.2 | 1,18E+04 | 1,18E+04 | 99.73 | 4570 | 31 | 1,18E+04 | 1,18E+04 |
| BRZ-4_S4_L001_R1_001 | MG649975 | Jul 24, 2016 | PA 3.2 | 23.91 | 17.36 | ViralCapSeq-VERT | 1,72E+06 | 1,67E+06 | 1,52E+05 | 9.09 | 1,17E+04 | NC_004162.2 | 1,18E+04 | 1,18E+04 | 99.74 | 1806 | 30 | 1,18E+04 | 1,18E+04 |
| BRZ-6_S5_L001_R1_001 | MG649983 | Mar 28, 2016 | PA 1.0 | 24.07 | 19.42 | ViralCapSeq-VERT | 2,22E+06 | 2,16E+06 | 6,74E+04 | 3.12 | 1,28E+04 | NC_004162.2 | 1,18E+04 | 1,18E+04 | 100 | 751 | NA | NA | NA |
| BRZ-7_S6_L001_R1_001 | MG649980 | Apr 26, 2016 | PA 1.0 | 25.20 | 16.30 | ViralCapSeq-VERT | 2,35E+06 | 2,28E+06 | 8,73E+05 | 38.29 | 1,17E+04 | NC_004162.2 | 1,18E+04 | 1,18E+04 | 99.94 | 10198 | 7 | 1,18E+04 | 1,18E+04 |
| BRZ-8_S7_L001_R1_001 | MG649979 | Mar 03, 2017 | PA 1.0 | 25.24 | 29.41 | ViralCapSeq-VERT | 2,66E+06 | 2,58E+06 | 3,29E+05 | 12.74 | 1,17E+04 | NC_004162.2 | 1,18E+04 | 1,18E+04 | 99.89 | 3880 | 13 | 1,18E+04 | 1,18E+04 |
| CHK_5_Brazil-RJ_2015 | MG649984 | 2015 | ND | ND | ND | Unbiased sequence | 2,15E+06 | 1,98E+06 | 1,93E+05 | 9.73 | 1,12E+04 | No reference | No reference | 1,20E+04 | 100 | 3790 | NA | NA | NA |
| CHK_7_Brazil-RJ_2015 | MG649985 | 2015 | ND | ND | ND | Unbiased sequence | 8,43E+05 | 7,64E+05 | 5,18E+02 | 0.07 | 1,12E+04 | No reference | No reference | 1,12E+04 | 95.02 | 19.72 | NA | NA | NA |

*Reference to Figure S3

*Legend:*

ND - Not determined (donated by Arbovirus Reference Laboratory and derived from their surveillance)

NA - Not applicable

**Table S3.** Chikungunya virus complete-genome sequences used for phylogenetic analyses.

| Acession Number | Analysis | Country | Host | Isolation source | Colection date | Passage history |
|---|---|---|---|---|---|---|
| KR559493 | ML | American Samoa | | | 2014 | |
| HM045823 | ML | Angola | | | 1962 | |
| KY435455 | ML | Anguilla | Homo sapiens | cell culture supernatant (vero) | 12/nov/14 | Passage in cell culture |
| KY435483 | ML | Anguilla | Homo sapiens | cell culture supernatant (vero) | 12-Feb-2014 | Passage in cell culture |
| KY435479 | ML | Antigua - Barbuda | Homo sapiens | cell culture supernatant (vero) | 28-Apr-2014 | Passage in cell culture |
| KY435470 | ML | Bahamas | Homo sapiens | cell culture supernatant (vero) | 08/jul/14 | Passage in cell culture |
| FJ807898 | ML | Bangladesh | Homo sapiens | infected patient | 2008 | |
| KU365370 | ML | Bangladesh | Homo sapiens | | nov/11 | |
| KU365371 | ML | Bangladesh | Homo sapiens | | nov/11 | |
| KY435464 | ML | Barbados | Homo sapiens | cell culture supernatant (vero) | 15-Aug-2014 | Passage in cell culture |
| KY435466 | ML | Barbados | Homo sapiens | cell culture supernatant (vero) | 06-Aug-2014 | Passage in cell culture |
| KP164568 | ML | Brazil | Homo sapiens | | 26-Aug-2014 | |
| KP164570 | ML | Brazil | Homo sapiens | | 03-Sep-2014 | |
| KP164571 | ML | Brazil | Homo sapiens | | 03/jul/14 | |
| KP164572 | ML | Brazil | Homo sapiens | | 21-Aug-2014 | |

| | | | | | |
|---|---|---|---|---|---|
| KT581 023 | ML | Brazil | Homo sapiens | | 2014 |
| KU355 832 | ML | Brazil | Homo sapiens | | 2015 |
| KU940 225 | ML | Brazil | Homo sapiens | | 15/jul /15 |
| KU940 226 | ML | Brazil | Homo sapiens | | 01- Aug- 2015 |
| KX228 391 | ML | Brazil | Homo sapiens | | 03/ma r/16 |
| KY055 011 | ML | Brazil | Aedes aegypti | | 20- Feb- 2016 |
| KY704 933 | ML /Ba yes | Brazil | Homo sapiens | | 30/ma r/16 |
| KY704 934 | ML /Ba yes | Brazil | Homo sapiens | | 30/ma r/16 |
| KY704 935 | ML /Ba yes | Brazil | Homo sapiens | | 01- Apr- 2016 |
| KY704 936 | ML /Ba yes | Brazil | Homo sapiens | | 01- Apr- 2016 |
| KY704 937 | ML /Ba yes | Brazil | Homo sapiens | | 15- Apr- 2016 |
| KY704 938 | ML /Ba yes | Brazil | Homo sapiens | | 19- Apr- 2016 |
| KY704 939 | ML /Ba yes | Brazil | Homo sapiens | | 17- Apr- 2016 |
| KY704 940 | ML /Ba yes | Brazil | Homo sapiens | | 14- Apr- 2016 |
| KY704 941 | ML /Ba yes | Brazil | Homo sapiens | | 14- Apr- 2016 |
| KY704 942 | ML /Ba yes | Brazil | Homo sapiens | | 16- Apr- 2016 |

| | | | | | |
|---|---|---|---|---|---|
| KY704 943 | ML /Ba yes | Brazil | Homo sapiens | | 19-Apr-2016 |
| KY704 944 | ML /Ba yes | Brazil | Homo sapiens | | 19-Apr-2016 |
| KY704 945 | ML /Ba yes | Brazil | Homo sapiens | | 14-Apr-2016 |
| KY704 946 | ML /Ba yes | Brazil | Homo sapiens | | 17-Apr-2016 |
| KY704 947 | ML /Ba yes | Brazil | Homo sapiens | | 15-Apr-2016 |
| KY704 948 | ML /Ba yes | Brazil | Homo sapiens | | 19-Apr-2016 |
| KY704 949 | ML /Ba yes | Brazil | Homo sapiens | | 07-Apr-2016 |
| KY704 950 | ML /Ba yes | Brazil | Homo sapiens | | 07-Apr-2016 |
| KY704 951 | ML /Ba yes | Brazil | Homo sapiens | | 13-Apr-2016 |
| KY704 952 | ML /Ba yes | Brazil | Homo sapiens | | 07-Apr-2016 |
| KY704 953 | ML /Ba yes | Brazil | Homo sapiens | | 09-Apr-2016 |
| KY704 954 | ML /Ba yes | Brazil | Homo sapiens | | 20/jun /16 |
| KY704 955 | ML /Ba yes | Brazil | Homo sapiens | | 17/jun /16 |
| KP164 567 | ML | Brazil-AP | Homo sapiens | | 28-Aug-2014 |
| KP164 569 | ML | Brazil-BA | Homo sapiens | | 28-Aug-2014 |

| | | | | | | |
|---|---|---|---|---|---|---|
| KY124328 | ML | Brazil-RJ | Homo sapiens | patient with febrile illness | 16/mar/16 | |
| KY124329 | ML | Brazil-RJ | Homo sapiens | patient with febrile illness | 16/mar/16 | |
| KJ451624 | ML | British Virgin | Homo sapiens | | jan/14 | |
| KY435486 | ML | British Virgin | Homo sapiens | cell culture supernatant (vero) | 23/jan/14 | Passage in cell culture |
| JQ861253 | ML | Cambodia | Homo sapiens | | 16-Aug-2011 | |
| JQ861254 | ML | Cambodia | Homo sapiens | | 16-Aug-2011 | |
| JQ861255 | ML | Cambodia | Homo sapiens | | 16-Aug-2011 | |
| JQ861256 | ML | Cambodia | Homo sapiens | | 16-Aug-2011 | |
| JQ861257 | ML | Cambodia | Homo sapiens | | 16-Aug-2011 | |
| JQ861258 | ML | Cambodia | Homo sapiens | | 16-Aug-2011 | |
| JQ861259 | ML | Cambodia | Homo sapiens | | 26-May-2011 | |
| JQ861260 | ML | Cambodia | Homo sapiens | | 28-May-2011 | |
| KX262996 | ML/Bayes | Cameroon | Homo sapiens | | 2006 | passage history: Vero 3; genotype: East-Central-South-African |
| HM045784 | ML/Bayes | CAR | Anopheles (Ceilia) funestus | | 19/nov/84 | |
| HM045822 | ML/Bayes | CAR | Homo sapiens | | Oct-1978 | |
| KY038946 | ML/Bayes | CAR | Aedes opok | | jun/75 | viral sample derived from a lyophilized vial |

| | | | | | | |
|---|---|---|---|---|---|---|
| KY038947 | ML/Bayes | CAR | Homo sapiens | | Dec-1983 | viral sample derived from a lyophilized vial; CRORAcollection |
| KY435459 | ML | Cayman-Islands | Homo sapiens | cell culture supernatant (vero) | 17-Sep-2014 | Passage in cell culture |
| KY435460 | ML | Cayman-Islands | Homo sapiens | cell culture supernatant (vero) | 06/jul/14 | Passage in cell culture |
| GU199350 | ML | China | Homo sapiens | serum | 2008 | |
| GU199351 | ML | China | Homo sapiens | serum | 2008 | passaged once in Vero cells |
| GU199352 | ML | China | Homo sapiens | serum | 2008 | |
| GU199353 | ML | China | Homo sapiens | serum | 2008 | |
| HQ846356 | ML | China | Homo sapiens | serum | Oct-2010 | |
| HQ846357 | ML | China | Homo sapiens | serum | Oct-2010 | |
| HQ846358 | ML | China | Homo sapiens | serum | Oct-2010 | |
| HQ846359 | ML | China | Homo sapiens | serum | Oct-2010 | |
| JQ065885 | ML | China | Homo sapiens | | Oct-2010 | |
| JQ065886 | ML | China | Homo sapiens | | Oct-2010 | |
| JQ065887 | ML | China | Homo sapiens | | Oct-2010 | |
| JQ065888 | ML | China | Homo sapiens | | Oct-2010 | |
| JQ065889 | ML | China | Homo sapiens | | Oct-2010 | |
| JQ065890 | ML | China | Homo sapiens | | Oct-2010 | |
| JQ065891 | ML | China | Homo sapiens | | Oct-2010 | |
| JQ065892 | ML | China | Homo sapiens | | Oct-2010 | |
| JX088705 | ML | China | Homo sapiens | | 2010 | |

| | | | | | | |
|---|---|---|---|---|---|---|
| KC488650 | ML | China | Homo sapiens | infected patient | 2012 | |
| KF318729 | ML | China | Homo sapiens | | 06/jul/12 | |
| KR559491 | ML | Colombia | | | Aug-2014 | |
| KU365372 | ML | Colombia | Homo sapiens | | 18-Dec-2014 | |
| KU365373 | ML | Colombia | Homo sapiens | | Dec-2014 | |
| KX496989 | ML | Colombia | Homo sapiens | blood | 09-Feb-2016 | |
| KF283986 | ML | Comoros | Homo sapiens | | 2005 | |
| KF283987 | ML | Comoros | Homo sapiens | | 2005 | |
| KP702297 | ML | Comoros | Homo sapiens | | 2005 | |
| KP003813 | ML/Bayes | Congo | Homo sapiens | | 2011 | |
| HM045818 | ML | Cote-dIvoire | Aedes luteocephalus | | Sep-1981 | |
| HM045820 | ML | Cote-dIvoire | Aedes africanus | | Dec-1993 | |
| KY435484 | ML | Dominica | Homo sapiens | cell culture supernatant (vero) | 30/jan/14 | Passage in cell culture |
| KY435485 | ML | Dominica | Homo sapiens | cell culture supernatant (vero) | 28/jan/14 | Passage in cell culture |
| KR559477 | ML | Dominican-Republic | | | jul/14 | |
| KR559479 | ML | Dominican-Republic | | | Apr-2014 | |
| KR559498 | ML | Dominican-Republic | | | mar/14 | |

| | | | | | |
|---|---|---|---|---|---|
| KY272961 | ML | Dominican-Republic | Homo sapiens | sera | 2014 |
| KY272962 | ML | Dominican-Republic | Homo sapiens | sera | 2014 |
| KY272963 | ML | Dominican-Republic | Homo sapiens | sera | 2014 |
| KY272964 | ML | Dominican-Republic | Homo sapiens | sera | 2014 |
| KY272965 | ML | Dominican-Republic | Homo sapiens | sera | 2014 |
| KY272966 | ML | Dominican-Republic | Homo sapiens | sera | 2014 |
| KY272967 | ML | Dominican-Republic | Homo sapiens | sera | 2014 |
| KY272968 | ML | Dominican-Republic | Homo sapiens | sera | 2014 |
| KY272969 | ML | Dominican-Republic | Homo sapiens | sera | 2014 |
| KY272970 | ML | Dominican-Republic | Homo sapiens | sera | 2014 |
| HM045809 | ML/Bayes | DRC | Homo sapiens | | 1960 |
| KR559471 | ML | El-Salvador | | | Oct-2014 |

| | | | | | | |
|---|---|---|---|---|---|---|
| KR559472 | ML | El-Salvador | | | jun/14 | |
| KR559475 | ML | El-Salvador | | | Sep-2014 | |
| KR559484 | ML | El-Salvador | | | nov/14 | |
| KP003807 | ML | France | Homo sapiens | | 2006 | |
| KR559473 | ML | French Polynesia | | | Feb-2015 | |
| KX262994 | ML | French-Guiana | Homo sapiens | | 21/jan/14 | passage history: C636#2, Vero 1 |
| KP003812 | ML/Bayes | Gabon | Homo sapiens | | 2007 | |
| KY435469 | ML | Grenada | Homo sapiens | cell culture supernatant (vero) | 30/jul/14 | Passage in cell culture |
| KY435472 | ML | Grenada | Homo sapiens | cell culture supernatant (vero) | 16/jun/14 | Passage in cell culture |
| KX262992 | ML | Guadeloupe | Homo sapiens | | 05/jan/14 | passage history: C636#2, Vero 1 |
| LN898094 | ML | Guadeloupe | Homo sapiens | | jan/14 | |
| LN898097 | ML | Guadeloupe | Homo sapiens | | jan/14 | |
| LN898098 | ML | Guadeloupe | Homo sapiens | | jan/14 | |
| LN898099 | ML | Guadeloupe | Homo sapiens | | jan/14 | |
| LN898102 | ML | Guadeloupe | Homo sapiens | | jan/14 | |
| LN898103 | ML | Guadeloupe | Homo sapiens | | jan/14 | |
| LN898110 | ML | Guadeloupe | Homo sapiens | | jan/14 | |
| LN898111 | ML | Guadeloupe | Homo sapiens | | jan/14 | |
| KR559481 | ML | Guatemala | | | Sep-2014 | |

| | | | | | | |
|---|---|---|---|---|---|---|
| KR559 490 | ML | Guyana | | | Aug-2014 | |
| KR559 496 | ML | Guyana | | | jul/14 | |
| KY435 458 | ML | Guyana | Homo sapiens | cell culture supernatant (vero) | 03/nov/14 | Passage in cell culture |
| KY435 477 | ML | Guyana | Homo sapiens | cell culture supernatant (vero) | 31-May-2014 | Passage in cell culture |
| KY435 478 | ML | Guyana | Homo sapiens | cell culture supernatant (vero) | 17-May-2014 | Passage in cell culture |
| KR559 476 | ML | Haiti | | | jul/14 | |
| KR559 478 | ML | Haiti | | | May-2014 | |
| KX702 401 | ML | Haiti | Homo sapiens | plasma | 02/jun/14 | |
| KX702 402 | ML | Haiti | Homo sapiens | plasma | 09/jun/14 | |
| KY415 978 | ML | Haiti | Homo sapiens | plasma | 29-May-2014 | |
| KY415 979 | ML | Haiti | Homo sapiens | plasma | 29-May-2014 | |
| KY415 980 | ML | Haiti | Homo sapiens | plasma | 05/jun/14 | |
| KY415 981 | ML | Haiti | Homo sapiens | plasma | 10/jun/14 | |
| KY415 982 | ML | Haiti | Homo sapiens | plasma | 11/jun/14 | |
| KY415 983 | ML | Haiti | Homo sapiens | plasma | 02/jun/14 | |
| KY415 984 | ML | Haiti | Homo sapiens | plasma | 24/jun/14 | |
| KY415 985 | ML | Haiti | Homo sapiens | plasma | 13-Aug-2014 | |
| KY435 480 | ML | Haiti | Homo sapiens | cell culture supernatant (vero) | 27-Apr-2014 | Passage in cell culture |

| | | | | | | |
|---|---|---|---|---|---|---|
| KR559487 | ML | Honduras | | | jul/14 | |
| KR559488 | ML | Honduras | | | Sep-2014 | |
| EF210157 | ML | India | Homo sapiens | | 2006 | |
| EU372006 | ML | India | Homo sapiens | | 11/jun/07 | |
| EU564335 | ML | India | Homo sapiens | supernatant of infected Vero B4 cells | 31-Oct-2006 | passaged in Vero B4 cells |
| FJ000062 | ML | India | Homo sapiens | CSF | Sep-2006 | |
| FJ000063 | ML | India | Homo sapiens | CSF | Oct-2006 | |
| FJ000064 | ML | India | Homo sapiens | CSF | Sep-2006 | |
| FJ000065 | ML | India | Homo sapiens | CSF | Sep-2006 | |
| FJ000066 | ML | India | Homo sapiens | serum | Sep-2006 | |
| FJ000067 | ML | India | Homo sapiens | CSF | Aug-2006 | |
| FJ000068 | ML | India | Homo sapiens | CSF | Aug-2006 | |
| FJ000069 | ML | India | Homo sapiens | serum | jun/07 | |
| GQ428210 | ML | India | Homo sapiens | serum | 07-Oct-2006 | |
| GQ428211 | ML | India | Homo sapiens | serum | 07-Oct-2006 | |
| GQ428212 | ML | India | Homo sapiens | serum | 12/jul/07 | |
| GQ428213 | ML | India | Homo sapiens | serum | 13/jul/07 | |
| GQ428214 | ML | India | Homo sapiens | serum | 29/jun/08 | |
| GQ428215 | ML | India | Homo sapiens | serum | 29-May-2008 | |

| | | | | | |
|---|---|---|---|---|---|
| HM045788 | ML | India | Homo sapiens | | 1973 |
| HM045803 | ML | India | Homo sapiens | | 06/nov/63 |
| HM045806 | ML | India | | | 1986 |
| HM045813 | ML | India | Homo sapiens | | 06/nov/63 |
| JF274082 | ML | India | Homo sapiens | serum | 27-Sep-2006 |
| JN558834 | ML | India | Homo sapiens | | 2009 |
| JN558835 | ML | India | Homo sapiens | | 2008 |
| JN558836 | ML | India | Homo sapiens | | 2009 |
| KJ679577 | ML | India | Homo sapiens | serum of a patient visiting Calcutta School of Tropical Medicine | 12-Sep-2011 |
| KJ679578 | ML | India | Homo sapiens | serum of a patient visiting Calcutta School of Tropical Medicine | 21-Dec-2011 |
| KJ796844 | ML | India | | serum | 04-Aug-2009 |
| KJ796845 | ML | India | | serum | 04-Aug-2009 |
| KJ796846 | ML | India | | serum | 18-Aug-2009 |
| KT336777 | ML | India | | human serum | 08-Apr-2009 |
| KT336778 | ML | India | | human serum | 16/jul/10 |
| KT336779 | ML | India | | human serum | 03-Aug-2012 |
| KT336780 | ML | India | | human serum | 03-Aug-2012 |

| | | | | | | |
|---|---|---|---|---|---|---|
| KT336781 | ML | India | | human serum | 06/mar/13 | |
| KT336782 | ML | India | | human serum | 27-May-2013 | |
| KX619424 | ML | India | Homo sapiens | serum | 22-Sep-2015 | |
| KX619425 | ML | India | Homo sapiens | serum | 11-Sep-2015 | |
| KX619426 | ML | India | Homo sapiens | serum | 03-Dec-2015 | |
| KY057363 | ML | India | Homo sapiens | | 28-Aug-2016 | |
| KY751908 | ML | India | male patient | | 2016 | |
| FJ807897 | ML | Indonesia | Homo sapiens | infected patient | 2007 | |
| HM045791 | ML | Indonesia | Homo sapiens | | 1983 | |
| HM045797 | ML | Indonesia | Homo sapiens | | 1985 | |
| KC862329 | ML | Indonesia | Homo sapiens | serum | 2010 | |
| KM673291 | ML | Indonesia | Homo sapiens | serum | jan/13 | |
| EU244823 | ML | Italy | | | 2007 | |
| KP003810 | ML | Italy | Homo sapiens | | 2007 | |
| KP003811 | ML | Italy | Homo sapiens | | 2007 | |
| KX262989 | ML | Italy | Homo sapiens | | 2007 | passage history: P1, Vero 1; genotype: East-Central-South-African |
| KX262993 | ML | Italy | Homo sapiens | | 2007 | passage history: P1(3308), Vero 1 |
| KR559489 | ML | Jamaica | | | Oct-2014 | |

| | | | | | | |
|---|---|---|---|---|---|---|
| KY435 461 | ML | Jamaica | Homo sapiens | cell culture supernatant (vero) | 24-Aug-2014 | Passage in cell culture |
| KY435 462 | ML | Jamaica | Homo sapiens | cell culture supernatant (vero) | 25-Aug-2014 | Passage in cell culture |
| KY435 468 | ML | Jamaica | Homo sapiens | cell culture supernatant (vero) | 06-Aug-2014 | Passage in cell culture |
| FR717 336 | ML | La-Reunion | Homo sapiens | host serum | 26-Dec-2005 | |
| FR717 337 | ML | La-Reunion | Homo sapiens | host cornea | 26-Dec-2005 | |
| KP003 808 | ML | Madaga scar | Homo sapiens | | 2006 | |
| EU703 759 | ML | Malaysi a | Homo sapiens | infected patient | 2006 | passaged twice in C636 cells |
| EU703 760 | ML | Malaysi a | Homo sapiens | infected patient | 2006 | passaged twice in C636 cells |
| EU703 761 | ML | Malaysi a | Homo sapiens | infected patient | 2006 | passaged twice in C636 cells |
| EU703 762 | ML | Malaysi a | Homo sapiens | infected patient | 2006 | passaged twice in C636 cells |
| FJ8078 99 | ML | Malaysi a | Homo sapiens | infected patient | 2008 | |
| FN295 483 | ML | Malaysi a | Homo sapiens | host serum | mar/0 6 | |
| FN295 484 | ML | Malaysi a | Homo sapiens | host serum | mar/0 6 | |
| FN295 485 | ML | Malaysi a | Homo sapiens | host serum | 2008 | |
| FN295 487 | ML | Malaysi a | Homo sapiens | host serum | 2008 | |
| FR687 340 | ML | Malaysi a | Homo sapiens | human serum | 21-Aug-2008 | |
| FR687 341 | ML | Malaysi a | Homo sapiens | human serum | 05/no v/08 | |
| FR687 342 | ML | Malaysi a | Homo sapiens | human serum | 10/no v/08 | |

| | | | | | | |
|---|---|---|---|---|---|---|
| FR687343 | ML | Malaysia | Homo sapiens | human serum | 12-Dec-2008 | |
| FR687344 | ML | Malaysia | Homo sapiens | human serum | 12/jan/09 | |
| FR687345 | ML | Malaysia | Homo sapiens | human serum | 05-Feb-2009 | |
| FR687346 | ML | Malaysia | Homo sapiens | human serum | 16-Feb-2009 | |
| FR687347 | ML | Malaysia | Homo sapiens | human serum | 12-Feb-2009 | |
| FR687348 | ML | Malaysia | Homo sapiens | human serum | 03-Apr-2009 | |
| KM923917 | ML | Malaysia | Macaca fascicularis | | 09/mar/07 | |
| KM923918 | ML | Malaysia | Macaca fascicularis | | 09/mar/07 | |
| KM923919 | ML | Malaysia | Macaca fascicularis | | 09/mar/07 | |
| KM923920 | ML | Malaysia | Macaca fascicularis | | 09/mar/07 | |
| KX168429 | ML | Malaysia | Homo sapiens | serum | 2009 | |
| KX262997 | ML | Malaysia | Homo sapiens | | 2009 | passage history: C636#1 |
| LN898093 | ML | Martinique | Homo sapiens | | Dec-2013 | |
| LN898095 | ML | Martinique | Homo sapiens | | jan/14 | |
| LN898096 | ML | Martinique | Homo sapiens | | jan/14 | |
| LN898100 | ML | Martinique | Homo sapiens | | jan/14 | |
| LN898101 | ML | Martinique | Homo sapiens | | jan/14 | |
| LN898104 | ML | Martinique | Homo sapiens | | jan/14 | |
| LN898105 | ML | Martinique | Homo sapiens | | jan/14 | |

| | | | | | | |
|---|---|---|---|---|---|---|
| LN898 106 | ML | Martiniq ue | Homo sapiens | | jan/14 | |
| LN898 107 | ML | Martiniq ue | Homo sapiens | | jan/14 | |
| LN898 108 | ML | Martiniq ue | Homo sapiens | | jan/14 | |
| LN898 109 | ML | Martiniq ue | Homo sapiens | | jan/14 | |
| LN898 112 | ML | Martiniq ue | Homo sapiens | | jan/14 | |
| EU564 334 | ML | Mauritiu s | Homo sapiens | supernatant of infected Vero B4 cells | 14-Feb-2006 | passaged in Vero B4 cells |
| FJ9591 03 | ML | Mauritiu s | Homo sapiens | | 2006 | |
| KP003 809 | ML | Mayotte | Homo sapiens | | 2006 | |
| KP851 709 | ML | Mexico | Homo sapiens; female | serum | 15-Oct-2014 | |
| KP851 710 | ML | Mexico | Homo sapiens; female | cell culture | 30-May-2014 | Passage in cell culture |
| KT327 163 | ML | Mexico | Homo sapiens | serum | 2014 | |
| KT327 164 | ML | Mexico | Homo sapiens | serum | 2014 | |
| KT327 165 | ML | Mexico | Homo sapiens | serum | 2014 | |
| KT327 166 | ML | Mexico | Homo sapiens | serum | 2014 | |
| KT327 167 | ML | Mexico | Homo sapiens | serum | 2014 | |
| KU365 366 | ML | Mexico | Homo sapiens | | 09-Oct-2014 | |
| KU365 367 | ML | Mexico | Homo sapiens | | 07/no v/14 | |
| KU365 368 | ML | Mexico | Homo sapiens | | 26/ma r/14 | |
| KJ4516 22 | ML | Microne sia | Homo sapiens | | Oct-2013 | |

| | | | | | | |
|---|---|---|---|---|---|---|
| KJ451623 | ML | Micronesia | Homo sapiens | | Oct-2013 | |
| KJ689452 | ML | Micronesia | | Aedes aegypti mosquito pool | nov/13 | |
| KJ689453 | ML | Micronesia | | Aedes hensilli mosquito pool | nov/13 | |
| KY435457 | ML | Montserrat | Homo sapiens | cell culture supernatant (vero) | 30-Oct-2014 | Passage in cell culture |
| KY435467 | ML | Montserrat | Homo sapiens | cell culture supernatant (vero) | 24/jul/14 | Passage in cell culture |
| KF151174 | ML | Myanmar | Homo sapiens | serum | 13/jul/09 | |
| KF151175 | ML | Myanmar | Homo sapiens | serum | 11-Dec-2009 | |
| KF590564 | ML | Myanmar | Homo sapiens | infected patient | 2010 | |
| KF590565 | ML | Myanmar | Homo sapiens | infected patient | 2010 | |
| KF590566 | ML | Myanmar | Homo sapiens | infected patient | 2010 | |
| KF590567 | ML | Myanmar | Homo sapiens | infected patient | 2010 | |
| HE806461 | ML | New-Caledonia | Homo sapiens | host serum | 28-Feb-2011 | |
| KT192707 | ML | Nicaragua | Homo sapiens | | 31-Oct-2014 | |
| KY703888 | ML | Nicaragua | Homo sapiens | | 15-Aug-2015 | passage history: not passaged; |
| KY703889 | ML | Nicaragua | Homo sapiens | | 05-Aug-2015 | passage history: not passaged; |
| KY703890 | ML | Nicaragua | Homo sapiens | | 23-Sep-2015 | passage history: not passaged; |
| KY703891 | ML | Nicaragua | Homo sapiens | | 17-Sep-2015 | passage history: not passaged; |
| KY703892 | ML | Nicaragua | Homo sapiens | | 26/nov/15 | passage history: not passaged; |

| | | | | | | |
|---|---|---|---|---|---|---|
| KY703 893 | ML | Nicarag ua | Homo sapiens | | 02-Sep-2015 | passage history: unknown; |
| KY703 894 | ML | Nicarag ua | Homo sapiens | | 09-Oct-2015 | passage history: not passaged; |
| KY703 895 | ML | Nicarag ua | Homo sapiens | | 26/nov/15 | passage history: not passaged; |
| KY703 896 | ML | Nicarag ua | Homo sapiens | | 18-Dec-2014 | passage history: passaged; |
| KY703 897 | ML | Nicarag ua | Homo sapiens | | 16/jan/15 | passage history: not passaged; |
| KY703 898 | ML | Nicarag ua | Homo sapiens | | 25-Aug-2015 | passage history: not passaged; |
| KY703 899 | ML | Nicarag ua | Homo sapiens | | 20-Oct-2015 | passage history: not passaged; |
| KY703 900 | ML | Nicarag ua | Homo sapiens | | 14-Dec-2015 | passage history: not passaged; |
| KY703 901 | ML | Nicarag ua | Homo sapiens | | 26/jan/15 | passage history: not passaged; |
| KY703 902 | ML | Nicarag ua | Homo sapiens | | 05-Sep-2015 | passage history: unknown; |
| KY703 903 | ML | Nicarag ua | Homo sapiens | | 05-Sep-2015 | passage history: not passaged; |
| KY703 904 | ML | Nicarag ua | Homo sapiens | | 04-Oct-2014 | passage history: passaged; |
| KY703 905 | ML | Nicarag ua | Homo sapiens | | 17-Sep-2015 | passage history: not passaged; |
| KY703 906 | ML | Nicarag ua | Homo sapiens | | 08-Sep-2015 | passage history: not passaged; |
| KY703 907 | ML | Nicarag ua | Homo sapiens | | 15/jul/15 | passage history: not passaged; |
| KY703 908 | ML | Nicarag ua | Homo sapiens | | 02-Dec-2014 | passage history: passaged; |

| | | | | | | |
|---|---|---|---|---|---|---|
| KY703 909 | ML | Nicarag ua | Homo sapiens | | 28/jan /15 | passage history: not passaged; |
| KY703 910 | ML | Nicarag ua | Homo sapiens | | 18- Oct- 2015 | passage history: not passaged; |
| KY703 911 | ML | Nicarag ua | Homo sapiens | | 17- Dec- 2015 | passage history: not passaged; |
| KY703 912 | ML | Nicarag ua | Homo sapiens | | 14- Dec- 2015 | passage history: not passaged; |
| KY703 913 | ML | Nicarag ua | Homo sapiens | | 13- Aug- 2015 | passage history: not passaged; |
| KY703 914 | ML | Nicarag ua | Homo sapiens | | 15- Aug- 2015 | passage history: not passaged; |
| KY703 915 | ML | Nicarag ua | Homo sapiens | | 03- Aug- 2015 | passage history: not passaged; |
| KY703 916 | ML | Nicarag ua | Homo sapiens | | 04- Aug- 2015 | passage history: not passaged; |
| KY703 917 | ML | Nicarag ua | Homo sapiens | | 08/jan /16 | passage history: not passaged; |
| KY703 918 | ML | Nicarag ua | Homo sapiens | | 26/jan /15 | passage history: not passaged; |
| KY703 919 | ML | Nicarag ua | Homo sapiens | | 29/jan /15 | passage history: not passaged; |
| KY703 920 | ML | Nicarag ua | Homo sapiens | | 16/no v/15 | passage history: not passaged; |
| KY703 921 | ML | Nicarag ua | Homo sapiens | | 04/no v/15 | passage history: not passaged; |
| KY703 922 | ML | Nicarag ua | Homo sapiens | | 06- Sep- 2015 | passage history: unknown; |
| KY703 923 | ML | Nicarag ua | Homo sapiens | | 19- Sep- 2015 | passage history: not passaged; |
| KY703 924 | ML | Nicarag ua | Homo sapiens | | 13- Aug- 2015 | passage history: not passaged; |

| | | | | | | |
|---|---|---|---|---|---|---|
| KY703925 | ML | Nicaragua | Homo sapiens | | 06-Sep-2015 | passage history: unknown; |
| KY703926 | ML | Nicaragua | Homo sapiens | | 20-Aug-2015 | passage history: not passaged; |
| KY703927 | ML | Nicaragua | Homo sapiens | | 15/jan/15 | passage history: not passaged; |
| KY703928 | ML | Nicaragua | Homo sapiens | | 30/jan/15 | passage history: not passaged; |
| KY703929 | ML | Nicaragua | Homo sapiens | | 02-Dec-2015 | passage history: not passaged; |
| KY703930 | ML | Nicaragua | Homo sapiens | | 24/nov/15 | passage history: not passaged; |
| KY703931 | ML | Nicaragua | Homo sapiens | | 19-Sep-2015 | passage history: not passaged; |
| KY703932 | ML | Nicaragua | Homo sapiens | | 19-Aug-2015 | passage history: not passaged; |
| KY703933 | ML | Nicaragua | Homo sapiens | | 09-Oct-2015 | passage history: not passaged; |
| KY703934 | ML | Nicaragua | Homo sapiens | | 27-Oct-2015 | passage history: not passaged; |
| KY703935 | ML | Nicaragua | Homo sapiens | | 02-Dec-2015 | passage history: not passaged; |
| KY703936 | ML | Nicaragua | Homo sapiens | | 07-Oct-2015 | passage history: not passaged; |
| KY703937 | ML | Nicaragua | Homo sapiens | | 15-Oct-2015 | passage history: not passaged; |
| HM045786 | ML | Nigeria | Homo sapiens | | 07/jul/64 | |
| HM045807 | ML | Nigeria | sentinel mouse | brain | 13-Apr-1965 | |
| KR559486 | ML | Panama | | | nov/14 | |

| | | | | | | |
|---|---|---|---|---|---|---|
| AB860301 | ML | Philippines | Homo sapiens | patient serum | 2013 | |
| HM045790 | ML | Philippines | Homo sapiens | | 17/jul/85 | |
| HM045800 | ML | Philippines | Homo sapiens | | 1985 | |
| KT308159 | ML | Philippines | Homo sapiens | serum | 2012 | |
| KT308160 | ML | Philippines | Homo sapiens | serum | 2012 | |
| KT308161 | ML | Philippines | Homo sapiens | serum | 2012 | |
| KT308162 | ML | Philippines | Homo sapiens | serum | 2012 | |
| KT308163 | ML | Philippines | Homo sapiens | serum | 2012 | |
| KR264949 | ML | Puerto-Rico | Homo sapiens | | 15/jul/14 | |
| KR264950 | ML | Puerto-Rico | Homo sapiens | | 16/jul/14 | |
| KR264951 | ML | Puerto-Rico | Homo sapiens | | 14-Aug-2014 | |
| KR559470 | ML | Puerto-Rico | | | nov/14 | |
| KR559474 | ML | Puerto-Rico | | | Sep-2014 | |
| KR559483 | ML | Puerto-Rico | | | Oct-2014 | |
| KR559495 | ML | Puerto-Rico | | | jul/14 | |
| KF872195 | ML | Russia | Homo sapiens | blood | 24-Sep-2013 | |
| KY435481 | ML | Saint | Homo sapiens | cell culture supernatant (vero) | 22-Apr-2014 | Passage in cell culture |
| KY435482 | ML | Saint | Homo sapiens | cell culture supernatant (vero) | 11/mar/14 | Passage in cell culture |
| KR559492 | ML | Saint Lucia | | | Aug-2014 | |

| | | | | | | |
|---|---|---|---|---|---|---|
| KY435474 | ML | Saint Lucia | Homo sapiens | cell culture supernatant (vero) | 19-May-2014 | Passage in cell culture |
| KX262991 | ML | Saint Martin | Homo sapiens | | 2003 | passage history: P2J3, Vero 2; |
| KY435473 | ML | Saint Vincent | Homo sapiens | cell culture supernatant (vero) | 22-May-2014 | Passage in cell culture |
| KY435475 | ML | Saint Vincent | Homo sapiens | cell culture supernatant (vero) | 22-May-2014 | Passage in cell culture |
| HM045785 | ML | Senegal | Aedes aegypti | | nov/66 | |
| HM045798 | ML | Senegal | Homo sapiens | | nov/66 | |
| HM045815 | ML | Senegal | Aedes luteocephalus | | Feb-1979 | |
| HM045816 | ML | Senegal | Homo sapiens | | 23/nov/66 | |
| HM045817 | ML | Senegal | Homo sapiens | | nov/05 | |
| HM045819 | ML | Senegal | Aedes dalzieli | | Feb-1993 | |
| HM045821 | ML/Bayes | Senegal | Chiroptera | | 20/mar/63 | |
| KX262986 | ML | Senegal | Aedes furcifer | | 1983 | passage history: AP61#1, Vero 2; |
| KX262995 | ML | Senegal | Aedes furcifer | | 10/jul/83 | passage history: AP61#1, Vero1; genotype: West African |
| FJ445430 | ML | Singapore | Homo sapiens | serum | jul/08 | |
| FJ445431 | ML | Singapore | Homo sapiens | serum | jul/08 | |
| FJ445432 | ML | Singapore | Homo sapiens | serum | jul/08 | |
| FJ445433 | ML | Singapore | Homo sapiens | serum | Aug-2008 | |
| FJ445443 | ML | Singapore | Homo sapiens | serum | Aug-2008 | |

| | | | | | | |
|---|---|---|---|---|---|---|
| FJ445445 | ML | Singapore | Homo sapiens | serum | Aug-2008 | |
| FJ445463 | ML | Singapore | Homo sapiens | serum | jul/08 | |
| FJ445484 | ML | Singapore | Homo sapiens | serum | May-2008 | |
| FJ445502 | ML | Singapore | Homo sapiens | serum | Aug-2008 | |
| FJ445510 | ML | Singapore | Homo sapiens | serum | jan/08 | |
| FJ445511 | ML | Singapore | Homo sapiens | serum | jan/08 | |
| FJ807896 | ML | Singapore | Homo sapiens | infected patient | 2006 | |
| HM045792 | ML/Bayes | South-Africa | Homo sapiens | | Apr-1956 | |
| HM045795 | ML/Bayes | South-Africa | Homo sapiens | | 1976 | |
| HM045805 | ML/Bayes | South-Africa | Aedes furcifer | | 1976 | |
| AB455493 | ML | Sri-Lanka | Homo sapiens | infected patient | 2006-12 | passaged in Vero B4 cells |
| AB455494 | ML | Sri-Lanka | Homo sapiens | infected patient | 2006-12 | passaged in Vero B4 cells |
| FJ445426 | ML | Sri-Lanka | Homo sapiens | serum | Apr-2008 | |
| FJ445427 | ML | Sri-Lanka | Homo sapiens | serum | jul/07 | |
| FJ445428 | ML | Sri-Lanka | Homo sapiens | serum | May-2007 | |
| FJ513628 | ML | Sri-Lanka | Homo sapiens | serum | mar/08 | |
| FJ513629 | ML | Sri-Lanka | Homo sapiens | serum | mar/08 | |
| FJ513632 | ML | Sri-Lanka | Homo sapiens | serum | mar/08 | |
| FJ513635 | ML | Sri-Lanka | Homo sapiens | serum | mar/08 | |

| | | | | | | |
|---|---|---|---|---|---|---|
| FJ513637 | ML | Sri-Lanka | Homo sapiens | serum | mar/08 | |
| FJ513645 | ML | Sri-Lanka | Homo sapiens | serum | Apr-2008 | |
| FJ513654 | ML | Sri-Lanka | Homo sapiens | serum | Apr-2008 | |
| FJ513657 | ML | Sri-Lanka | Homo sapiens | serum | Apr-2008 | |
| FJ513673 | ML | Sri-Lanka | Homo sapiens | serum | Apr-2008 | |
| FJ513675 | ML | Sri-Lanka | Homo sapiens | serum | Apr-2008 | |
| FJ513679 | ML | Sri-Lanka | Homo sapiens | serum | Apr-2008 | |
| GU013528 | ML | Sri-Lanka | Homo sapiens | serum | mar/08 | |
| GU013529 | ML | Sri-Lanka | Homo sapiens | serum | mar/08 | |
| GU013530 | ML | Sri-Lanka | Homo sapiens | serum | Apr-2008 | |
| GU189061 | ML | Sri-Lanka | Homo sapiens | | 2006 | |
| HM045799 | ML | Sri-Lanka | Homo sapiens | | 2007 | |
| HM045801 | ML | Sri-Lanka | Homo sapiens | | 2007 | |
| KR559497 | ML | St. Barts | | | jun/14 | |
| KY435456 | ML | Suriname | Homo sapiens | cell culture supernatant (vero) | 17-Aug-2014 | Passage in cell culture |
| KY435463 | ML | Suriname | Homo sapiens | cell culture supernatant (vero) | 02-Aug-2014 | Passage in cell culture |
| HM045811 | ML/Bayes | Tanzania | Homo sapiens | | 22-Feb-1953 | |
| GQ905863 | ML | Thailand | Homo sapiens | | 25-May-2009 | |
| GU301779 | ML | Thailand | Homo sapiens | serum | 04-Sep-2009 | |

| | | | | | | |
|---|---|---|---|---|---|---|
| GU301780 | ML | Thailand | Homo sapiens | serum | 21-Oct-2008 | |
| GU301781 | ML | Thailand | Homo sapiens | culture | 27/jul/09 | Passage in cell culture |
| GU908223 | ML | Thailand | Aedes albopictus | supernatant | 14-Aug-2009 | Passage in cell culture |
| HM045787 | ML | Thailand | Homo sapiens | | 1995 | |
| HM045789 | ML | Thailand | Homo sapiens | | 1988 | |
| HM045796 | ML | Thailand | Homo sapiens | | 1995 | |
| HM045808 | ML | Thailand | Homo sapiens | | 1978 | |
| HM045810 | ML | Thailand | Homo sapiens | | 1958 | |
| HM045814 | ML | Thailand | Homo sapiens | | 1975 | |
| KJ579184 | ML | Thailand | Homo sapiens | serum sample from suspected patient | 14-Oct-2013 | |
| KJ579185 | ML | Thailand | Homo sapiens | serum sample from suspected patient | 14-Oct-2013 | |
| KJ579186 | ML | Thailand | Homo sapiens | serum sample from suspected patient | 14-Oct-2013 | |
| KJ579187 | ML | Thailand | Homo sapiens | serum sample from suspected patient | 14-Oct-2013 | |
| KJ796847 | ML | Thailand | | serum | 05/mar/09 | |
| KJ796848 | ML | Thailand | | age 57 female serum | 06/nov/08 | |
| KJ796849 | ML | Thailand | | age 28 male serum | 23-Dec-2008 | |
| KJ796850 | ML | Thailand | | age 48 female serum | 16-Sep-2009 | |

| | | | | | | |
|---|---|---|---|---|---|---|
| KJ796851 | ML | Thailand | | age 60 female serum | 01-Dec-2008 | |
| KJ796852 | ML | Thailand | | age 56 female serum | 12-Feb-2009 | |
| KP164869 | ML | Thailand | | | 2009 | |
| KX009167 | ML | Thailand | Homo sapiens | serum | 2013 | |
| KX009168 | ML | Thailand | Homo sapiens | serum | 2013 | |
| KX009169 | ML | Thailand | Homo sapiens | serum | 2013 | |
| KX009170 | ML | Thailand | Homo sapiens | serum | 2013 | |
| KX009171 | ML | Thailand | Homo sapiens | serum | 2013 | |
| KX262987 | ML | Thailand | Homo sapiens | | 1996 | passage history: LLC-MK#2, Vero 1 |
| KX262988 | ML | Thailand | Homo sapiens | | 1988 | passage history: LLC-MK2#2,C636#1, Vero 1; |
| KR046227 | ML | Trinidad | Homo sapiens | serum | 30-Aug-2014 | |
| KR046228 | ML | Trinidad | Homo sapiens | serum | 11-Sep-2014 | |
| KR046229 | ML | Trinidad | Homo sapiens | serum | 17-Sep-2014 | |
| KR046230 | ML | Trinidad | Homo sapiens | serum | 18-Sep-2014 | |
| KR046231 | ML | Trinidad | Homo sapiens | serum | 09/nov/14 | |
| KR046232 | ML | Trinidad | Homo sapiens | serum | 20-Sep-2014 | |
| KR046233 | ML | Trinidad | Homo sapiens | serum | 20-Sep-2014 | |

| | | | | | | |
|---|---|---|---|---|---|---|
| KR046234 | ML | Trinidad | Homo sapiens | serum | 12-Sep-2014 | |
| KU365369 | ML | Trinidad | Homo sapiens | | 2014 | |
| KY435454 | ML | Trinidad | Homo sapiens | cell culture supernatant (vero) | nov/11 | Passage in cell culture |
| KY435465 | ML | Trinidad | Homo sapiens | cell culture supernatant (vero) | 17-Aug-2014 | Passage in cell culture |
| KY435471 | ML | Turks and Caicos | Homo sapiens | cell culture supernatant (vero) | 11/jun/14 | Passage in cell culture |
| KY435476 | ML | Turks and Caicos | Homo sapiens | cell culture supernatant (vero) | 05/jun/14 | Passage in cell culture |
| HM045812 | ML/Bayes | Uganda | Homo sapiens | | 1982 | |
| HM045794 | ML | USA | Homo sapiens | | 2006 | |
| KJ941050 | ML | USA | Homo sapiens | cell culture supernatant, Vero passage 4 | 2006 | Passage in cell culture |
| KY575565 | ML | USA | Homo sapiens | | 2014 | passage history: V1; |
| KY575566 | ML | USA | Homo sapiens | | 2014 | passage history: V1; |
| KY575567 | ML | USA | Homo sapiens | | 2006 | passage history: P?V2 |
| KY575568 | ML | USA | Homo sapiens | | 2006 | passage history: P?V2 |
| KY575569 | ML | USA | Homo sapiens | | 2014 | passage history: V1; |
| KY575570 | ML | USA | Homo sapiens | | 2008 | passage history: V1 |
| KY575571 | ML | USA | Homo sapiens | | 2006 | passage history: V2 |
| KY575572 | ML | USA | Homo sapiens | | 2014 | passage history: V1; |
| KY575573 | ML | USA | Homo sapiens | | 2014 | passage history: V1; |
| KY575574 | ML | USA | Homo sapiens | | 1995 | passage history: P?SM1V3 |

| | | | | | | |
|---|---|---|---|---|---|---|
| KY680 347 | ML | USA | Homo sapiens | | 13-May-2014 | passage history: VERO 0; |
| KY680 348 | ML | USA | Homo sapiens | | 05-Sep-2014 | |
| KY680 349 | ML | USA | Homo sapiens | | 02/jul /14 | passage history: VERO 0; |
| KY680 350 | ML | USA | Homo sapiens | | 10-Dec-2014 | |
| KY680 351 | ML | USA | Homo sapiens | | 24/jun /14 | passage history: VERO 0; |
| KY680 352 | ML | USA | Homo sapiens | | 15-May-2014 | |
| KY680 353 | ML | USA | Homo sapiens | | 25-Aug-2014 | |
| KY680 354 | ML | USA | Homo sapiens | | 08-May-2014 | passage history: VERO 0; |
| KY680 355 | ML | USA | Homo sapiens | | 22/jul /14 | |
| KY680 356 | ML | USA | Homo sapiens | | 21/jul /14 | passage history: VERO 0; |
| KY680 357 | ML | USA | Homo sapiens | | 09-Aug-2014 | passage history: VERO 0; |
| KY680 358 | ML | USA | Homo sapiens | | 08-Aug-2014 | passage history: VERO 0; |
| KY680 359 | ML | USA | Homo sapiens | | 04/jul /14 | passage history: VERO 0; |
| KY680 360 | ML | USA | Homo sapiens | | 05-Aug-2014 | |
| KY680 361 | ML | USA | Homo sapiens | | 26-May-2014 | |
| KY680 362 | ML | USA | Homo sapiens | | 26/jul /14 | passage history: VERO 0; |
| KY680 363 | ML | USA | Homo sapiens | | 15/jun /14 | passage history: VERO 0; |

| | | | | | | |
|---|---|---|---|---|---|---|
| KY680 364 | ML | USA | Homo sapiens | | 15-May-2014 | |
| KY680 365 | ML | USA | Homo sapiens | | 18-Sep-2014 | |
| KY680 366 | ML | USA | Homo sapiens | | 24/jul /15 | |
| KY680 367 | ML | USA | Homo sapiens | | 09/jun /14 | passage history: VERO 0; |
| KY680 368 | ML | USA | Homo sapiens | | 01-Dec-2014 | |
| KY680 369 | ML | USA | Homo sapiens | | 24-May-2014 | passage history: VERO 0; |
| KY680 370 | ML | USA | Homo sapiens | | 03-Oct-2014 | |
| KY680 371 | ML | USA | Homo sapiens | | 20-Aug-2014 | passage history: VERO 0; |
| KY680 372 | ML | USA | Homo sapiens | | 04/jun /14 | passage history: VERO 0; |
| KY680 373 | ML | USA | Homo sapiens | | 22-Oct-2014 | |
| KY680 374 | ML | USA | Homo sapiens | | 06-Dec-2014 | |
| KY680 375 | ML | USA | Homo sapiens | | 04/jun /14 | passage history: VERO 0; |
| KY680 376 | ML | USA | Homo sapiens | | 08-Oct-2014 | |
| KY680 377 | ML | USA | Homo sapiens | | 16-Aug-2014 | |
| KY680 378 | ML | USA | Homo sapiens | | 19-Oct-2014 | |
| KY680 379 | ML | USA | Homo sapiens | | 16-Sep-2014 | |

| | | | | | | |
|---|---|---|---|---|---|---|
| KY680 380 | ML | USA | Homo sapiens | | 13-Oct-2014 | |
| KY680 381 | ML | USA | Homo sapiens | | 21-Oct-2014 | |
| KY680 382 | ML | USA | Homo sapiens | | 04/jun/14 | |
| KY680 383 | ML | USA | Homo sapiens | | 20-Aug-2014 | |
| KY680 384 | ML | USA | Homo sapiens | | 25/jun/14 | passage history: VERO 0; |
| KY680 385 | ML | USA | Homo sapiens | | 25/nov/14 | |
| KY680 386 | ML | USA | Homo sapiens | | 24-Sep-2014 | |
| KY680 387 | ML | USA | Homo sapiens | | 14-Oct-2014 | |
| KY680 388 | ML | USA | Homo sapiens | | 02-Oct-2014 | |
| KY680 389 | ML | USA | Homo sapiens | | 14/jul/15 | |
| KY680 390 | ML | USA | Homo sapiens | | 07-May-2014 | passage history: VERO 0; |
| KY680 391 | ML | USA | Homo sapiens | | 24-Oct-2014 | |
| KY680 392 | ML | USA | Homo sapiens | | 19-Oct-2014 | |
| KY680 393 | ML | USA | Homo sapiens | | 17-Sep-2014 | |
| KY680 394 | ML | USA | Homo sapiens | | 05-Oct-2014 | |
| KY680 395 | ML | USA | Homo sapiens | | 21/jul/14 | passage history: VERO 0; |

| | | | | | | |
|---|---|---|---|---|---|---|
| KY680 396 | ML | USA | Homo sapiens | | 02-Sep-2014 | |
| KY680 397 | ML | USA | Homo sapiens | | 23-May-2014 | passage history: VERO 0; |
| KY680 398 | ML | USA | Homo sapiens | | 05-Aug-2014 | |
| KY680 399 | ML | USA | Homo sapiens | | 05-Sep-2014 | |
| KY680 400 | ML | USA | Homo sapiens | | 15/jul/14 | passage history: VERO 0; |
| KY680 401 | ML | USA | Homo sapiens | | 22-Sep-2014 | |
| KY680 402 | ML | USA | Homo sapiens | | 27/jul/14 | passage history: VERO 0; |
| KY680 403 | ML | USA | Homo sapiens | | 14/jun/14 | passage history: VERO 0; |
| KY680 404 | ML | USA | Homo sapiens | | 17-Sep-2014 | |
| KY680 405 | ML | USA | Homo sapiens | | 18/jun/14 | passage history: VERO 0; |
| KY680 406 | ML | USA | Homo sapiens | | 08-Aug-2014 | passage history: VERO 0; |
| KY680 407 | ML | USA | Homo sapiens | | 20/nov/14 | |
| KY680 408 | ML | USA | Homo sapiens | | 26/jun/14 | passage history: VERO 0; |
| KY680 409 | ML | USA | Homo sapiens | | 21-Oct-2014 | |
| KY680 410 | ML | USA | Homo sapiens | | 22-Aug-2014 | passage history: VERO 0; |
| KY680 411 | ML | USA | Homo sapiens | | 04-Sep-2014 | |

| | | | | | | |
|---|---|---|---|---|---|---|
| KY680412 | ML | USA | Homo sapiens | | 28-Aug-2014 | |
| KY680413 | ML | USA | Homo sapiens | | 17-Sep-2014 | |
| KY680414 | ML | USA | Homo sapiens | | 13-Sep-2014 | |
| MF001507 | ML | USA | Homo sapiens | | 2015 | passaged 1 time in C636 |
| MF001508 | ML | USA | Homo sapiens | | 2015 | passaged 1 time in C636 |
| MF001509 | ML | USA | Homo sapiens | | 2015 | passaged 1 time in C636 |
| MF001510 | ML | USA | Homo sapiens | | 2015 | passaged 1 time in C636 |
| MF001511 | ML | USA | Homo sapiens | | 2015 | passaged 1 time in C636 |
| MF001512 | ML | USA | Homo sapiens | | 2015 | passaged 1 time in C636 |
| MF001513 | ML | USA | Homo sapiens | | 2015 | passaged 1 time in C636 |
| MF001514 | ML | USA | Homo sapiens | | 2015 | passaged 1 time in C636 |
| MF001515 | ML | USA | Homo sapiens | | 2015 | passaged 1 time in C636 |
| MF001516 | ML | USA | Homo sapiens | | 2015 | passaged 1 time in C636 |
| MF001517 | ML | USA | Homo sapiens | | 2015 | passaged 1 time in C636 |
| MF001518 | ML | USA | Homo sapiens | | 2015 | passaged 1 time in C636 |
| MF001519 | ML | USA | Homo sapiens | | 2015 | passaged 1 time in C636 |
| KU365374 | ML | Venezuela | Homo sapiens | | 17-Dec-2014 | |
| KR559480 | ML | Virgin-Islands | | | Sep-2014 | |
| KR559482 | ML | Virgin-Islands | | | Aug-2014 | |

| | | | | | |
|---|---|---|---|---|---|
| KR559 485 | ML | Virgin-Islands | | | Oct-2014 |
| KR559 494 | ML | Virgin-Islands | | | jul/14 |
| KC614 648 | ML | Yemen | Aedes aegypti | | 25/jan /11 |

ML: Maximum-likelihood; Bayes: Bayesian phylogeographic; USA: United States of America; CAR: Central African Republic. Empty spaces mean that these information are not public available.

**Table S4 - Socio-Demographical data from Patients tested by RT-PCR**

| Year | 2016 | 2017 |
|---|---|---|
| ***Number of individual tested by RT-PCR (n)*** | 827 | 999 |
| ***Number of individual with detectable viral loads (n)*** | 233 | 189 |
| ***Detectable molecular diagnosis (%)*** | | |
| CHIKV | 72 | 37 |
| DENV-4 | 12 | 17 |
| ZIKV | 16 | 47 |
| ***Gender (%)*** | | |
| Female | 58 | 57 |
| ***Age [median years-old (IQR)]*** | 34 (23-45) | 30 (21-45) |
| ***Rio de Janeiro's subregion (%, Figure S3 for reference)*** | | |
| 1.0 | 34 | 8 |
| 2.1 | 8 | 12 |
| 2.2 | 5 | 7 |
| 3.1 | 2 | 4 |
| 3.2 | 13 | 14 |
| 3.3 | 18 | 18 |
| 4.0 | 10 | 13 |
| 5.1 | 4 | 8 |
| 5.2 | 3 | 14 |
| 5.3 | 3 | 2 |

**Table S5.** Best fit spatial, molecular clock and demographic models for the ECSA Bayesian phylogeographic analysis.

| Spatial | Clock | Demographic | PS | SS | Spatial | | | Clock | | | Demographic | | |
|---|---|---|---|---|---|---|---|---|---|---|---|---|---|
| | | | | | Models | PS BF | SS BF | Models | PS BF | SS BF | Models | PS BF | SS BF |
| Sym | Strict | Skygrid | -23174,82 | -23174,77 | Sym/Assym | -3,46 | 14,47 | Strict/UCLN | -6,14 | -6,24 | - | - | - |
| | | Skyline | -23152,65 | -23121,70 | | **9,93** | **40,96** | | **8,33** | **39,45** | Skyl/Skyg | **22,17** | **53,07** |
| | UCLN | Skygrid | -23168,68 | -23168,53 | | 0,42 | 0,46 | - | - | - | - | - | - |
| | | Skyline | -23160,98 | -23161,15 | | 2,15 | 2,37 | | - | - | Skyl/Skyg | 7,70 | 7,38 |
| Assym | Strict | Skygrid | -23171,36 | -23171,37 | - | - | - | Strict/UCLN | -2,26 | -2,39 | - | - | - |
| | | Skyline | -23162,58 | -23162,67 | - | - | - | | 0,55 | 0,85 | Skyl/Skyg | 8,78 | 8,71 |
| | UCLN | Skygrid | -23169,10 | -23168,99 | - | - | - | - | - | - | - | - | - |
| | | Skyline | -23163,13 | -23163,52 | - | - | - | | - | - | Skyl/Skyg | 5,97 | 5,47 |

Log marginal likelihood (ML) estimates for the phylogeographic models [symmetric (Sym) and asymmetric (Assym)], molecular-clock models [strict and uncorrelated log-normal (UCLN)], and non-parametric demographic models [Skygrid (Skyg) and Skyline (Skyl)] obtained using the path sampling (PS) and stepping-stone sampling (SS) methods. The Log Bayes factor (BF) is the difference of the Log ML between of alternative (H1) and null (H0) models (H1/H0). Log positive BFs values indicates that model H1 is more strongly supported by the data than model H0, while negative values indicate that BF is in favor of the model H0. Each model was systematically compared, and higher BF values were highlighted in bold while the best fit model was underlined.

### Table S6 - Polymorphisms in the CHIKV genome

| Mature peptide | Polymorphism Ty | Product | Nt Start | Nt Stop | CDS Type | CDS Positio | CDS Codon Numb | Protein position | Codon pos | Change | Polymorphism Ty | Codon Chang | Protein Effe | Amino Acid Chang | Variant Sequences |
|---|---|---|---|---|---|---|---|---|---|---|---|---|---|---|---|
| **Nonstructural protein nsP1 peptide** | | | 1 | 1.605 | | | 1 | 1 | | | | | | | |
| | SNP (transition) | nsP1 | 189 | | nonstructural | 189 | 63 | 63 | 3 | T -> C | SNP (transition) | GAT -> GAC | None | | ECSA BR ROOT |
| | SNP (transition) | nsP1 | 267 | | nonstructural | 267 | 89 | 89 | 3 | T -> C | SNP (transition) | GAT -> GAC | None | | ECSA AL NODE |
| | SNP (transition) | nsP1 | 309 | | nonstructural | 309 | 103 | 103 | 3 | C -> T | SNP (transition) | GCC -> GCT | None | | ECSA BR ROOT |
| | SNP (transition) | nsP1 | 405 | | nonstructural | 405 | 135 | 135 | 3 | A -> G | SNP (transition) | TTA -> TTG | None | | ECSA BR ROOT |
| | SNP (transition) | nsP1 | 603 | | nonstructural | 603 | 201 | 201 | 3 | T -> C | SNP (transition) | GAT -> GAC | None | | ECSA BR ROOT |
| | SNP (transition) | nsP1 | 669 | | nonstructural | 669 | 223 | 223 | 3 | C -> T | SNP (transition) | GGC -> GGT | None | | ECSA BR ROOT |
| | SNP (transition) | nsP1 | 708 | | nonstructural | 708 | 236 | 236 | 3 | C -> T | SNP (transition) | TGC -> TGT | None | | ECSA BR ROOT |
| | SNP (transition) | nsP1 | 945 | | nonstructural | 945 | 315 | 315 | 3 | T -> C | SNP (transition) | TGT -> TGC | None | | ECSA BR ROOT |
| | SNP (transition) | nsP1 | 1.389 | | nonstructural | 1.389 | 463 | 463 | 3 | T -> C | SNP (transition) | CCT -> CCC | None | | ECSA BR ROOT |
| | SNP (transition) | nsP1 | 1.449 | | nonstructural | 1.449 | 483 | 483 | 3 | C -> T | SNP (transition) | TAC -> TAT | None | | ECSA BR ROOT |
| **Non-structural protein nsP2 peptide; NTPase peptide** | | | 1.606 | 3.999 | | | 536 | | | | | | | | |
| | SNP (transversion) | nsP2 | 1.623 | | nonstructural | 1.623 | 541 | 6 | 3 | G -> T | SNP (transversion) | CCG -> CCT | None | | ECSA BR ROOT |
| | SNP (transition) | nsP2 | 1.785 | | nonstructural | 1.785 | 595 | 60 | 3 | T -> C | SNP (transition) | TAT -> TAC | None | | ECSA BR ROOT |
| | SNP (transition) | nsP2 | 1.920 | | nonstructural | 1.920 | 640 | 105 | 3 | T -> C | SNP (transition) | CAT -> CAC | None | | ECSA BR ROOT |
| | SNP (transition) | nsP2 | 2.169 | | nonstructural | 2.169 | 723 | 188 | 3 | G -> A | SNP (transition) | CCG -> CCA | None | | ECSA BR ROOT |
| | SNP (transition) | nsP2 | 2.337 | | nonstructural | 2.337 | 779 | 244 | 3 | A -> G | SNP (transition) | AGA -> AGG | None | | ECSA BR ROOT |
| | SNP (transition) | nsP2 | 2.436 | | nonstructural | 2.436 | 812 | 277 | 3 | T -> C | SNP (transition) | CTT -> CTC | None | | ECSA BR ROOT |
| | **SNP (transversion)** | **nsP2** | **2.659** | | **nonstructural** | **2.659** | **887** | **352** | **1** | **C -> G** | **SNP (transversion)** | **CCT -> GCT** | **Substitution** | **P -> A** | **ECSA AL NODE, ECSA RJ-SE NODE** |
| | SNP (transition) | nsP2 | 2.667 | | nonstructural | 2.667 | 889 | 354 | 3 | C -> T | SNP (transition) | GAC -> GAT | None | | ECSA BR ROOT |
| | SNP (transition) | nsP2 | 2.778 | | nonstructural | 2.778 | 926 | 391 | 3 | C -> T | SNP (transition) | TAC -> TAT | None | | ECSA BR ROOT |
| | SNP (transition) | nsP2 | 2.889 | | nonstructural | 2.889 | 963 | 428 | 3 | T -> C | SNP (transition) | GGT -> GGC | None | | ECSA AL NODE |
| | SNP (transition) | nsP2 | 2.946 | | nonstructural | 2.946 | 982 | 447 | 3 | T -> C | SNP (transition) | ATT -> ATC | None | | ECSA BR ROOT |
| | SNP (transition) | nsP2 | 3.033 | | nonstructural | 3.033 | 1.011 | 476 | 3 | C -> T | SNP (transition) | AAC -> AAT | None | | ECSA BR ROOT |
| | SNP (transition) | nsP2 | 3.132 | | nonstructural | 3.132 | 1.044 | 509 | 3 | C -> T | SNP (transition) | GAC -> GAT | None | | ECSA BR ROOT |
| | SNP (transition) | nsP2 | 3.193 | | nonstructural | 3.193 | 1.065 | 530 | 1 | C -> T | SNP (transition) | CTA -> TTA | None | | ECSA BR ROOT |
| | SNP (transition) | nsP2 | 3.201 | | nonstructural | 3.201 | 1.067 | 532 | 3 | C -> T | SNP (transition) | AGC -> AGT | None | | ECSA BR ROOT |
| | SNP (transition) | nsP2 | 3.219 | | nonstructural | 3.219 | 1.073 | 538 | 3 | G -> A | SNP (transition) | CCG -> CCA | None | | ECSA BR ROOT |
| | SNP (transition) | nsP2 | 3.285 | | nonstructural | 3.285 | 1.095 | 560 | 3 | T -> C | SNP (transition) | TTT -> TTC | None | | ECSA BR ROOT |
| | SNP (transition) | nsP2 | 3.369 | | nonstructural | 3.369 | 1.123 | 588 | 3 | T -> C | SNP (transition) | ACT -> ACC | None | | ECSA BR ROOT |
| | SNP (transition) | nsP2 | 3.405 | | nonstructural | 3.405 | 1.135 | 600 | 3 | C -> T | SNP (transition) | AAC -> AAT | None | | ECSA BR ROOT |
| | SNP (transition) | nsP2 | 3.420 | | nonstructural | 3.420 | 1.140 | 605 | 3 | C -> T | SNP (transition) | AAC -> AAT | None | | ECSA BR ROOT |
| | SNP (transition) | nsP2 | 3.534 | | nonstructural | 3.534 | 1.178 | 643 | 3 | C -> T | SNP (transition) | AAC -> AAT | None | | ECSA BR ROOT |
| | SNP (transition) | nsP2 | 3.539 | | nonstructural | 3.539 | 1.180 | 645 | 2 | C -> T | SNP (transition) | GCA -> GTA | Substitution | A -> V | ECSA BR ROOT |
| | SNP (transition) | nsP2 | 3.636 | | nonstructural | 3.636 | 1.212 | 677 | 3 | T -> C | SNP (transition) | GGT -> GGC | None | | ECSA BR ROOT |
| | SNP (transition) | nsP2 | 3.825 | | nonstructural | 3.825 | 1.275 | 740 | 3 | A -> G | SNP (transition) | GTA -> GTG | None | | ECSA BR ROOT |
| | SNP (transition) | nsP2 | 3.840 | | nonstructural | 3.840 | 1.280 | 745 | 3 | T -> C | SNP (transition) | TTT -> TTC | None | | ECSA BR ROOT |
| | SNP (transition) | nsP2 | 3.951 | | nonstructural | 3.951 | 1.317 | 782 | 3 | C -> T | SNP (transition) | AAC -> AAT | None | | ECSA BR ROOT |
| **Nonstructural protein nsP3 peptide** | | | 4.000 | 5.589 | | | 1334 | | | | | | | | |
| | SNP (transition) | nsP3 | 4.059 | | nonstructural | 4.059 | 1.353 | 20 | 3 | C -> T | SNP (transition) | GTC -> GTT | None | | ECSA BR ROOT |
| | SNP (transition) | nsP3 | 4.173 | | nonstructural | 4.173 | 1.391 | 58 | 3 | T -> C | SNP (transition) | GTT -> GTC | None | | ECSA BR ROOT |
| | SNP (transition) | nsP3 | 4.212 | | nonstructural | 4.212 | 1.404 | 71 | 3 | A -> G | SNP (transition) | CCA -> CCG | None | | ECSA BR ROOT |
| | SNP (transition) | nsP3 | 4.215 | | nonstructural | 4.215 | 1.405 | 72 | 3 | C -> T | SNP (transition) | AAC -> AAT | None | | ECSA RJ-SE NODE |
| | SNP (transition) | nsP3 | 4.229 | | nonstructural | 4.229 | 1.410 | 77 | 2 | C -> T | SNP (transition) | TCG -> TTG | Substitution | S -> L | ECSA BR ROOT |
| | SNP (transition) | nsP3 | 4.233 | | nonstructural | 4.233 | 1.411 | 78 | 3 | G -> A | SNP (transition) | GAG -> GAA | None | | ECSA BR ROOT |
| | SNP (transition) | nsP3 | 4.356 | | nonstructural | 4.356 | 1.452 | 119 | 3 | C -> T | SNP (transition) | GAC -> GAT | None | | ECSA BR ROOT |
| | SNP (transition) | nsP3 | 4.386 | | nonstructural | 4.386 | 1.462 | 129 | 3 | T -> C | SNP (transition) | TTT -> TTC | None | | ECSA BR ROOT |
| | SNP (transition) | nsP3 | 4.428 | | nonstructural | 4.428 | 1.476 | 143 | 3 | C -> T | SNP (transition) | TGC -> TGT | None | | ECSA BR ROOT |
| | SNP (transition) | nsP3 | 4.489 | | nonstructural | 4.489 | 1.497 | 164 | 1 | C -> T | SNP (transition) | CTG -> TTG | None | | ECSA BR ROOT |
| | SNP (transition) | nsP3 | 4.599 | | nonstructural | 4.599 | 1.533 | 200 | 3 | T -> C | SNP (transition) | TAT -> TAC | None | | ECSA AL NODE |
| | SNP (transition) | nsP3 | 4.611 | | nonstructural | 4.611 | 1.537 | 204 | 3 | C -> T | SNP (transition) | ACC -> ACT | None | | ECSA BR ROOT |
| | SNP (transition) | nsP3 | 4.635 | | nonstructural | 4.635 | 1.545 | 212 | 3 | T -> C | SNP (transition) | GAT -> GAC | None | | ECSA BR ROOT |
| | SNP (transition) | nsP3 | 4.641 | | nonstructural | 4.641 | 1.547 | 214 | 3 | G -> A | SNP (transition) | GCG -> GCA | None | | ECSA BR ROOT |
| | SNP (transition) | nsP3 | 4.761 | | nonstructural | 4.761 | 1.587 | 254 | 3 | A -> G | SNP (transition) | TCA -> TCG | None | | ECSA BR ROOT |
| | SNP (transition) | nsP3 | 4.767 | | nonstructural | 4.767 | 1.589 | 256 | 3 | C -> T | SNP (transition) | CCC -> CCT | None | | ECSA BR ROOT |
| | SNP (transversion) | nsP3 | 4.770 | | nonstructural | 4.770 | 1.590 | 257 | 3 | C -> A | SNP (transversion) | CCC -> CCA | None | | ECSA BR ROOT |
| | SNP (transition) | nsP3 | 4.809 | | nonstructural | 4.809 | 1.603 | 270 | 3 | A -> G | SNP (transition) | CCA -> CCG | None | | ECSA BR ROOT |
| | SNP (transition) | nsP3 | 4.848 | | nonstructural | 4.848 | 1.616 | 283 | 3 | C -> T | SNP (transition) | AGC -> AGT | None | | ECSA BR ROOT |
| | SNP (transition) | nsP3 | 4.863 | | nonstructural | 4.863 | 1.621 | 288 | 3 | T -> C | SNP (transition) | TCT -> TCC | None | | ECSA BR ROOT |
| | **SNP (transition)** | **nsP3** | **4.950** | | **nonstructural** | **4.950** | **1.650** | **317** | **3** | **G -> A** | **SNP (transition)** | **TCG -> TCA** | **None** | | **ECSA BR ROOT, ECSA RJ-SE NODE** |

| | | | | | | | | | | | | | | | |
|---|---|---|---|---|---|---|---|---|---|---|---|---|---|---|---|
| | SNP (transversion) | nsP3 | 4.987 | | nonstructural | 4.987 | 1.663 | 330 | 1 | T -> G | SNP (transversion) | TCT -> GCT | Substitution | S -> A | ECSA BR ROOT |
| | SNP (transition) | nsP3 | 5.061 | | nonstructural | 5.061 | 1.687 | 354 | 3 | A -> G | SNP (transition) | CTA -> CTG | None | | ECSA AL NODE |
| | SNP (transition) | nsP3 | 5.109 | | nonstructural | 5.109 | 1.703 | 370 | 3 | C -> T | SNP (transition) | GCC -> GCT | None | | ECSA BR ROOT |
| | SNP (transition) | nsP3 | 5.119 | | nonstructural | 5.119 | 1.707 | 374 | 1 | G -> A | SNP (transition) | GGG -> AGG | Substitution | G -> R | ECSA BR ROOT |
| | SNP (transition) | nsP3 | 5.148 | | nonstructural | 5.148 | 1.716 | 383 | 3 | T -> C | SNP (transition) | ACT -> ACC | None | | ECSA BR ROOT |
| | SNP (transition) | nsP3 | 5.345 | | nonstructural | 5.345 | 1.782 | 449 | 2 | C -> T | SNP (transition) | ACG -> ATG | Substitution | T -> M | ECSA BR ROOT |
| **Nonstructural protein nsP4 peptide** | | | 5.590 | 7.422 | | | 1864 | | | | | | | | |
| | SNP (transition) | nsP4 | 5.707 | | nonstructural | 5.707 | 1.903 | 40 | 1 | C -> T | SNP (transition) | CTG -> TTG | None | | ECSA BR ROOT |
| | SNP (transition) | nsP4 | 5.717 | | nonstructural | 5.717 | 1.906 | 43 | 2 | C -> T | SNP (transition) | GCA -> GTA | Substitution | A -> V | ECSA BR ROOT |
| | SNP (transition) | nsP4 | 5.866 | | nonstructural | 5.866 | 1.956 | 93 | 1 | C -> T | SNP (transition) | CCA -> TCA | Substitution | P -> S | ECSA BR ROOT |
| | SNP (transition) | nsP4 | 5.920 | | nonstructural | 5.920 | 1.974 | 111 | 1 | A -> G | SNP (transition) | ATC -> GTC | Substitution | I -> V | ECSA RJ NODE |
| | SNP (transition) | nsP4 | 6.034 | | nonstructural | 6.034 | 2.012 | 149 | 1 | C -> T | SNP (transition) | CTA -> TTA | None | | ECSA BR ROOT |
| | SNP (transition) | nsP4 | 6.126 | | nonstructural | 6.126 | 2.042 | 179 | 3 | C -> T | SNP (transition) | CAC -> CAT | None | | ECSA BR ROOT |
| | SNP (transition) | nsP4 | 6.273 | | nonstructural | 6.273 | 2.091 | 228 | 3 | C -> T | SNP (transition) | TTC -> TTT | None | | ECSA BR ROOT |
| | SNP (transition) | nsP4 | 6.396 | | nonstructural | 6.396 | 2.132 | 269 | 3 | C -> T | SNP (transition) | TTC -> TTT | None | | ECSA BR ROOT |
| | SNP (transition) | nsP4 | 6.408 | | nonstructural | 6.408 | 2.136 | 273 | 3 | T -> C | SNP (transition) | CAT -> CAC | None | | ECSA BR ROOT |
| | SNP (transition) | nsP4 | 6.630 | | nonstructural | 6.630 | 2.210 | 347 | 3 | C -> T | SNP (transition) | GAC -> GAT | None | | ECSA BR ROOT |
| | SNP (transition) | nsP4 | 6.717 | | nonstructural | 6.717 | 2.239 | 376 | 3 | T -> C | SNP (transition) | GAT -> GAC | None | | ECSA BR ROOT |
| | SNP (transition) | nsP4 | 6.750 | | nonstructural | 6.750 | 2.250 | 387 | 3 | T -> C | SNP (transition) | GCT -> GCC | None | | ECSA BR ROOT |
| | SNP (transition) | nsP4 | 6.757 | | nonstructural | 6.757 | 2.253 | 390 | 1 | C -> T | SNP (transition) | CTG -> TTG | None | | ECSA BR ROOT |
| | SNP (transition) | nsP4 | 6.813 | | nonstructural | 6.813 | 2.271 | 408 | 3 | C -> T | SNP (transition) | TTC -> TTT | None | | ECSA BR ROOT |
| | SNP (transversion) | nsP4 | 6.936 | | nonstructural | 6.936 | 2.312 | 449 | 3 | A -> C | SNP (transversion) | CGA -> CGC | None | | ECSA BR ROOT |
| | SNP (transversion) | nsP4 | 7.031 | | nonstructural | 7.031 | 2.344 | 481 | 2 | C -> A | SNP (transversion) | GCC -> GAC | Substitution | A -> D | ECSA RJ-SE NODE |
| | SNP (transition) | C protein | 7.146 | | nonstructural | 7.146 | 2.382 | 519 | 3 | T -> C | SNP (transition) | GCT -> GCC | None | | ECSA BR ROOT |
| **C protein peptide** | | | 7.491 | 8.273 | | 1 | 1 | 1 | | | | | | | |
| | SNP (transversion) | C protein | 7.498 | | structural | 8 | 3 | 3 | 2 | T -> A | SNP (transversion) | TTC -> TAC | Substitution | F -> Y | ECSA BR ROOT |
| | SNP (transition) | C protein | 7.559 | | structural | 69 | 23 | 23 | 3 | T -> C | SNP (transition) | CCT -> CCC | None | | ECSA BR ROOT |
| | SNP (transversion) | C protein | 7.571 | | structural | 81 | 27 | 27 | 3 | C -> A | SNP (transversion) | GTC -> GTA | None | | ECSA BR ROOT |
| | SNP (transition) | C protein | 7.604 | | structural | 114 | 38 | 38 | 3 | T -> C | SNP (transition) | GCT -> GCC | None | | ECSA BR ROOT |
| | SNP (transition) | C protein | 7.711 | | structural | 221 | 74 | 74 | 2 | A -> G | SNP (transition) | CAA -> CGA | Substitution | Q -> R | ECSA BR ROOT |
| | SNP (transition) | C protein | 7.835 | | structural | 345 | 115 | 115 | 3 | C -> T | SNP (transition) | TTC -> TTT | None | | ECSA BR ROOT |
| | SNP (transition) | C protein | 7.845 | | structural | 355 | 119 | 119 | 1 | C -> T | SNP (transition) | CAT -> TAT | Substitution | H -> Y | ECSA BR ROOT |
| | SNP (transition) | C protein | 8.039 | | structural | 549 | 183 | 183 | 3 | G -> A | SNP (transition) | CCG -> CCA | None | | ECSA BR ROOT |
| **E3 protein peptide** | | | 8.274 | 8.465 | | | 262 | | | | | | | | |
| | SNP (transition) | E3 | 8.301 | | structural | 811 | 271 | 10 | 1 | T -> C | SNP (transition) | TTG -> CTG | None | | ECSA BR ROOT |
| | SNP (transition) | E3 | 8.309 | | structural | 819 | 273 | 12 | 3 | C -> T | SNP (transition) | AAC -> AAT | None | | ECSA BR ROOT |
| | SNP (transversion) | E3 | 8.342 | | structural | 852 | 284 | 23 | 3 | A -> T | SNP (transversion) | ACA -> ACT | None | | ECSA BR ROOT |
| | SNP (transition) | E3 | 8.354 | | structural | 864 | 288 | 27 | 3 | C -> T | SNP (transition) | TAC -> TAT | None | | ECSA BR ROOT |
| | SNP (transition) | E3 | 8.359 | | structural | 869 | 290 | 29 | 2 | A -> G | SNP (transition) | AAG -> AGG | Substitution | K -> R | ECSA BR ROOT |
| | SNP (transition) | E3 | 8.456 | | structural | 966 | 322 | 61 | 3 | C -> T | SNP (transition) | CGC -> CGT | None | | ECSA BR ROOT |
| **E2 protein peptide** | | | 8.466 | 9.734 | | | 326 | | | | | | | | |
| | SNP (transition) | E2 | 8.507 | | structural | 1.017 | 339 | 14 | 3 | A -> G | SNP (transition) | CCA -> CCG | None | | ECSA BR ROOT |
| | SNP (transition) | E2 | 8.565 | | structural | 1.075 | 359 | 34 | 1 | C -> T | SNP (transition) | CTA -> TTA | None | | ECSA BR ROOT |
| | SNP (transition) | E2 | 8.686 | | structural | 1.196 | 399 | 74 | 2 | T -> C | SNP (transition) | ATG -> ACG | Substitution | M -> T | ECSA BR ROOT |
| | SNP (transition) | E2 | 8.693 | | structural | 1.203 | 401 | 76 | 3 | A -> G | SNP (transition) | GCA -> GCG | None | | ECSA BR ROOT |
| | SNP (transition) | E2 | 8.772 | | structural | 1.282 | 428 | 103 | 1 | G -> A | SNP (transition) | GCC -> ACC | Substitution | A -> T | ECSA BR ROOT |
| | SNP (transition) | E2 | 8.858 | | structural | 1.368 | 456 | 131 | 3 | C -> T | SNP (transition) | CAC -> CAT | None | | ECSA AL NODE |
| | SNP (transition) | E2 | 8.882 | | structural | 1.392 | 464 | 139 | 3 | A -> G | SNP (transition) | GAA -> GAG | None | | ECSA BR ROOT |
| | SNP (transition) | E2 | 8.918 | | structural | 1.428 | 476 | 151 | 3 | A -> G | SNP (transition) | CTA -> CTG | None | | ECSA BR ROOT |
| | SNP (transition) | E2 | 9.041 | | structural | 1.551 | 517 | 192 | 3 | T -> C | SNP (transition) | GTT -> GTC | None | | ECSA AL NODE |
| | SNP (transition) | E2 | 9.128 | | structural | 1.638 | 546 | 221 | 3 | G -> A | SNP (transition) | AAG -> AAA | None | | ECSA BR ROOT |
| | SNP (transition) | E2 | 9.176 | | structural | 1.686 | 562 | 237 | 3 | T -> C | SNP (transition) | TAT -> TAC | None | | ECSA BR ROOT |
| | SNP (transition) | E2 | 9.230 | | structural | 1.740 | 580 | 255 | 3 | T -> C | SNP (transition) | ATT -> ATC | None | | ECSA BR ROOT |
| | SNP (transition) | E2 | 9.332 | | structural | 1.842 | 614 | 289 | 3 | T -> C | SNP (transition) | CCT -> CCC | None | | ECSA BR ROOT |
| | SNP (transition) | E2 | 9.362 | | structural | 1.872 | 624 | 299 | 3 | T -> C | SNP (transition) | AAT -> AAC | None | | ECSA BR ROOT |
| | SNP (transition) | E2 | 9.383 | | structural | 1.893 | 631 | 306 | 3 | T -> C | SNP (transition) | TAT -> TAC | None | | ECSA BR ROOT |
| | SNP (transition) | E2 | 9.458 | | structural | 1.968 | 656 | 331 | 3 | C -> T | SNP (transition) | GGC -> GGT | None | | ECSA BR ROOT |
| **6K protein peptide** | | | 9.735 | 9.917 | | | 749 | | | | | | | | |
| | SNP (transition) | 6k | 9.770 | | structural | 2.280 | 760 | 12 | 3 | C -> T | SNP (transition) | AAC -> AAT | None | | ECSA BR ROOT |
| | SNP (transition) | 6k | 9.773 | | structural | 2.283 | 761 | 13 | 3 | G -> A | SNP (transition) | GAG -> GAA | None | | ECSA RJ NODE |
| | SNP (transition) | 6k | 9.860 | | structural | 2.370 | 790 | 42 | 3 | T -> C | SNP (transition) | TGT -> TGC | None | | ECSA BR ROOT |
| **E1 protein peptide** | | | 9.918 | 11.234 | | | 810 | | | | | | | | |
| | SNP (transition) | E1 | 10.079 | | structural | 2.589 | 863 | 54 | 3 | C -> T | SNP (transition) | GTC -> GTT | None | | ECSA BR ROOT |

| | | | | | | | | | | | | | | | |
|---|---|---|---|---|---|---|---|---|---|---|---|---|---|---|---|
| | SNP (transversion) | E1 | 10.190 | | structural | 2.700 | 900 | 91 | 3 | C -> A | SNP (transversion) | GGC -> GGA | None | | ECSA BR ROOT |
| | SNP (transition) | E1 | 10.238 | | structural | 2.748 | 916 | 107 | 3 | T -> C | SNP (transition) | CAT -> CAC | None | | ECSA BR ROOT |
| | SNP (transversion) | E1 | 10.361 | | structural | 2.871 | 957 | 148 | 3 | A -> T | SNP (transversion) | GCA -> GCT | None | | ECSA BR ROOT |
| | SNP (transversion) | E1 | 10.549 | | structural | 3.059 | 1.020 | 211 | 2 | A -> C | SNP (transversion) | AAA -> ACA | Substitution | K -> T | ECSA RJ-SE NODE |
| | SNP (transition) | E1 | 10.727 | | structural | 3.237 | 1.079 | 270 | 3 | C -> T | SNP (transition) | AAC -> AAT | None | | ECSA BR ROOT |
| | SNP (transition) | E1 | 10.780 | | structural | 3.290 | 1.097 | 288 | 2 | C -> T | SNP (transition) | ACT -> ATT | Substitution | T -> I | ECSA BR ROOT |
| | SNP (transversion) | E1 | 10.931 | | structural | 3.441 | 1.147 | 338 | 3 | T -> A | SNP (transversion) | ACT -> ACA | None | | ECSA BR ROOT |
| | SNP (transition) | E1 | 10.997 | | structural | 3.507 | 1.169 | 360 | 3 | A -> G | SNP (transition) | TTA -> TTG | None | | ECSA BR ROOT |
| | SNP (transition) | E1 | 11.047 | | structural | 3.557 | 1.186 | 377 | 2 | C -> T | SNP (transition) | GCA -> GTA | Substitution | A -> V | ECSA BR ROOT |
| | SNP (transversion) | E1 | 11.136 | | structural | 3.646 | 1.216 | 407 | 1 | A -> C | SNP (transversion) | ATG -> CTG | Substitution | M -> L | ECSA BR ROOT |
| | SNP (transition) | E1 | 11.165 | | structural | 3.675 | 1.225 | 416 | 3 | T -> C | SNP (transition) | GGT -> GGC | None | | ECSA BR ROOT |
| | SNP (transition) | E1 | 11.225 | | structural | 3.735 | 1.245 | 436 | 3 | T -> C | SNP (transition) | TTT -> TTC | None | | ECSA BR ROOT |
